# Induction of the ISR by AB5 subtilase cytotoxin drives type-I IFN expression in pDCs via STING activation

**DOI:** 10.1101/2024.10.07.616986

**Authors:** Daniela Barros, Beatriz H. Ferreira, Paulina Garcia-Gonzalez, Francesco Carbone, Marine Luka, Fátima Leite-Pinheiro, Paulo Antas, Andreia Mendes, Mariana D. Machado, Lichen Zhang, Marina Cresci, Lou Galliot, Julien Gigan, Marisa Reverendo, Bing Su, Adrienne W. Paton, James C. Paton, Stéphane Rocchi, Frédéric Rieux-Laucat, Rafael J. Argüello, Béatrice Nal, Yinming Liang, Mickaël Ménager, Evelina Gatti, Catarina R. Almeida, Philippe Pierre

## Abstract

We demonstrate that exposure to the AB5 subtilase cytotoxin (SubAB) induces the unfolded protein response (UPR) in human peripheral blood mononuclear cells, concomitant with a pro-inflammatory response across distinct cell subsets. Notably, SubAB selectively induces type-I interferon (IFN) expression in plasmacytoid dendritic cells, acting synergistically with Toll-like receptor 7 stimulation. The induction of type-I IFN in response to SubAB relies on stimulator of interferon genes (STING) activation, coupled with protein synthesis inhibition mediated by protein kinase R-like endoplasmic reticulum kinase and phosphorylation of the eukaryotic translation initiation factor 2 subunit-alpha. By impeding mRNA translation through the integrated stress response, SubAB precipitates the downregulation of the negative innate signaling feedback regulator Tax1-binding protein 1. This downregulation is necessary to unleash TANK-binding kinase 1 signaling associated with STING activation. These findings shed new light on how UPR-inducing conditions may regulate the immune system during infection or pathogenesis.

## INTRODUCTION

Environmental cues or infection can alter protein folding and flux through the endoplasmic reticulum (ER). Upon misfolded protein accumulation in the ER, several signaling pathways, known as the unfolded protein response (UPR), are induced to overcome biochemical stress and prevent premature cell death^1^. The UPR corrects excessive or pathological protein misfolding within the ER, by upregulating genes required for protein folding, glycosylation or degradation, as well as lipid production^2,3^. It is composed of three signaling branches, each triggered by the dissociation of the immunoglobulin-heavy-chain binding protein (BiP; also known as HSPA5/GRP78) from the ER-resident sensors, activating transcription factor 6 (ATF6), inositol-requiring enzyme 1-alpha (IRE1α) and protein kinase R (PKR)-like endoplasmic reticulum kinase (PERK or EIF2AK3)^4^. BiP dissociation from PERK in the ER lumen, induces its dimerization and activation to mediate eukaryotic initiation factor 2-alpha (eIF2α) phosphorylation in the cytosol. Phosphorylation of eIF2α on serine 51 converts eIF2 into an inhibitor of the GDP-GTP guanine exchange factor eIF2B, leading to a global reduction in protein synthesis, while favoring the translation of specific transcripts such as activating transcription factor 4 (ATF4) and protein phosphatase 1 regulatory subunit 15A (PPP1R15a/GADD34) to initiate the integrated stress response (ISR)^5,6^.

Plasmacytoid DCs (pDCs) and plasma cells, capable of producing large quantities of interferon (IFN) or immunoglobulins, respectively, are characterized by large ER volumes and display a dependence on IRE1α and spliced X-box-binding protein 1 (XBP1_s_) for their differentiation and function^7,8,9,10^. pDCs are a major source of type-I IFN in response to viral infections, which can become pathogenic during auto-immune disease, mainly through engagement of endosomal Toll-like receptor (TLR)7 and TLR9 by nucleic acids^11^. Although viral infection can trigger ER-stress, direct connection to type-I IFN expression remains largely uncertain, since ER stress inducers usually do not independently result in significant IFN production^12,3^. Alternatively, the cyclic dinucleotide c-GAMP is produced by the cytoplasmic DNA-sensing receptor cyclic GMP–AMP synthase (cGAS) to activate the ER-associated molecule stimulator of interferon genes (STING, TMEM173), and orchestrate anti-viral responses via TANK-binding kinase 1 (TBK1)- and interferon regulatory factor 3 (IRF3)-dependent type-I IFN production^13,14^. Cell-autonomous responses involving autophagy and direct PERK induction have been suggested to protect cell integrity from the co-lateral damages linked to STING activation^15,16,17^.

The subtilase cytotoxin (SubAB) was discovered in a Shiga toxigenic *Escherichia coli* strain (STEC) belonging to serotype O113:H21 that caused an outbreak of haemolytic uraemic syndrome (HUS) in South Australia^18^. SubAB belongs to the AB5 toxin family with the pentameric B subunit allowing cellular uptake and retro-translocation to the ER via its specificity for glycans terminating in N-glycolylneuraminic acid (Neu5Gc)^19,20,21^. The A subunit is a subtilase-like serine protease with exquisite substrate specificity for BiP in the ER^22^. The cleavage of BiP causes severe ER stress and an unresolved UPR in different cell types, ultimately resulting in their apoptosis^22^. Although its actual role in HUS is yet to be established, SubAB may have subtle effects on pathogenesis through immune modulation^23,24,25^, underlining the importance of the UPR in regulating inflammation^26,10^.

To unravel if SubAB could activate human immune cells and gain mechanistic insights on how ER stress could shape immunity, we analyzed the consequences of toxin exposure on primary human peripheral blood mononuclear cells (PBMCs) by single-cell RNA (scRNA) sequencing and spectral flow cytometry analysis. A clear signature of UPR induction was found in most cells analyzed, accompanied by an activation of nuclear factor kappa B (NF-kB)-associated pro-inflammatory pathways. Among PBMCs, non-classical CD16^+^ monocytes are the most sensitive to SubAB exposure. Additionally, the toxin synergizes with TLR7 signal to increase NF-kB-dependent signaling along type-I IFN responses in most cell subsets examined. We further show that SubAB drives type-I IFN production in primary blood pDCs, this also in strong synergy with TLR7 stimulation. At the molecular level, BiP inactivation results in type-I IFN expression via a combination of STING activation and concomitant targeted loss of negative innate signalling regulators, like Tax1-binding protein 1 (TAX1BP1). This sequence of events is completely dependent on PERK-mediated translation inhibition, and the capacity of SubAB to promote an overshooting anti-viral response in pDCs could represent a physiological mechanism by which HUS or IFN-dependent pathologies could be initiated in susceptible individuals^27^.

## RESULTS

### SubAB exposure drives the UPR and inflammatory responses in PBMCs

To comprehensively study the impact of SubAB on human cells, we performed multiparametric population analysis by flow cytometry and single-cell transcriptomics analysis of human PBMCs submitted to different stimulatory conditions, including 6 h of SubAB (1μg/ml) exposure and/or TLR7 stimulation with 1μM of the adenine analog CL307^28^. Uniform Manifold Approximation and Projection (UMAP) dimension reduction applied to multiparameter flow cytometry analysis of CD45^+^ cells from three healthy donors were performed (Fig. 1A). Following stimulation with SubAB, most identified cell types exhibited minimal alterations both in phenotype and number (Fig. 1B and Supplementary Fig. S1A). The toxin induced limited cell death, as monitored by absence of ViaDye Red^TM^ staining across most analyzed subsets (Supplementary Fig. S1B), validating our experimental approach and confirming that globally, like in mouse^23^, human immune cells are relatively resistant to ER stress-induced death. A heightened sensitivity of non-classical (CD14^dim^ CD16^+^) and intermediate (CD14^+^CD16^+^) monocytes^29^ to SubAB exposure was however observed, with a reduced number of these rare subsets compared to the conventional HLA-DR^+^CD14^+^CD16^-^ population (Fig.1C and Supplementary Fig. S1C).

**Figure 1.**
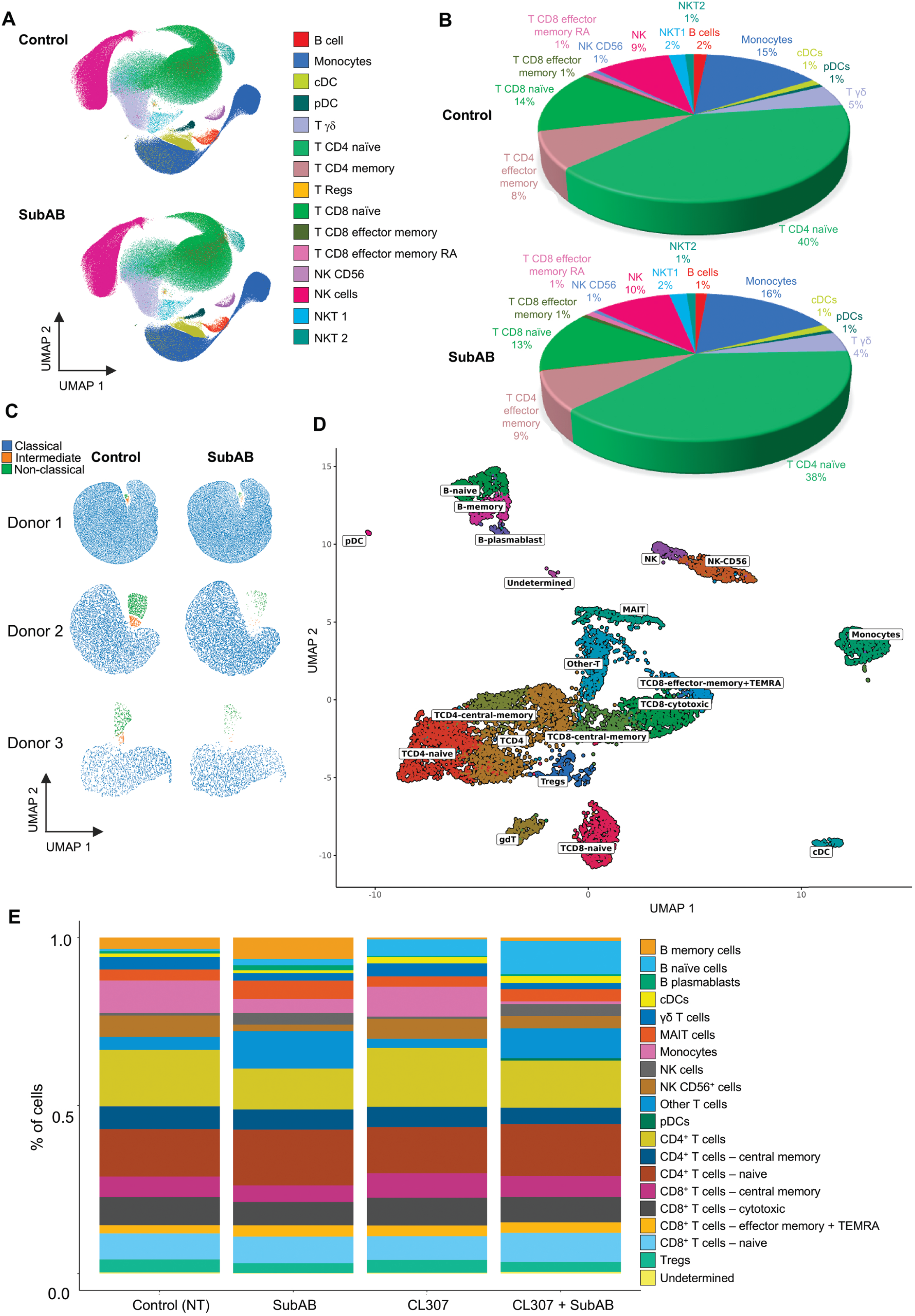
Multiparametric flow cytometry and scRNA seq analysis of human PBMCs stimulated with SubAB or CL307. **(A)** UMAP reduction and cell type assignment after multiparametric flow analysis of CD45^+^ PBMCs from 3 blood donors treated or not with 1 μg/ml of SubAB for 6 h. **(B)** Pie graph representation of the cell subsets proportions found by multiparametric flow analysis in PBMC treated (bottom) or not (top) with SubAB. Proportions were found very similar across conditions. **(C)** UMAP reduction of spectral flow analysis of monocytes after 6 h of SubAB treatment clustered on expression levels of CD14 and/or CD16. **(D)** UMAP and cell type assignment of 22449 PBMC from total scRNA seq dataset. **(E)** Boxplot of cell subsets proportion found across scRNA seq experimental conditions including Control, SubAB-treated, CL307-treated and CL307/SubAB-treated PBMCs. See also Sup Fig S1.

Next, by following rigorous quality control, integration, and unsupervised clustering of scRNA-seq data, we successfully identified 19 clusters representing most of the cell populations normally detected in blood PBMCs (Fig. 1D)^30^. As observed during our cytometry analysis, the proportions of most populations remain relatively stable upon SubAB exposure, except for the monocyte and natural killer (NK) CD56^+^ subsets, which were reduced (Fig. 1E and Supplementary Fig. S2A). CL307 also triggered some changes in the proportion of the main TLR7 expressing cells detected, mainly affecting the memory B cell subset, while rare pDCs were only detected in samples treated with both CL307 and SubAB, rendering their analysis across different conditions impossible (Fig. 1E and Supplementary Fig. S2A).

Comparative analysis of differentially expressed genes (Differential Expression Genes (DEGs) and Gene Ontology (GO) pathways) using the Enrichir bioinformatic suite^31^ in SubAB- and/or CL307-treated PBMCs versus untreated control cells confirmed that both treatments impacted gene expression with various efficiency in cell subsets (Supplementary Fig. S2B). As expected, SubAB most strongly impacted mRNA transcription in monocytes, with memory B cells, CD4^+^ and CD8^+^ T cells coming next. Clear induction of the UPR was confirmed in monocytes and B cells, as well as in NK and more modestly in DCs and T cell subsets (Fig. 2A and 2B and Supplementary Table S1). In monocytes, SubAB had also a clear transcriptional down-modulating effect (Supplementary Fig. S2B). Genes known to be preferentially expressed by CD16^+^ monocytes (e.g. *LYZ, IFI30, MTSS1, CXCL16, CD93, MAFB, TAGLN2, MS4A7,* and *LILRB1*)^29,32^ were down-regulated (Fig. 2C). This likely reflects the reduction in CD16^+^ non-classical and intermediate monocytes, which appear more sensitive to the toxin than the CD14^+^CD16^-^ classical population, as suggested by the comparative analysis of CD14 and CD16 expression in the different experimental conditions (Supplementary Fig. S2C). Despite the relative loss of CD14 expression in response to SubAB in the scRNA data set, classical monocytes still exhibited increased mRNA expression of the cytokine genes (e.g*. IL23a, IL6, IL18, Ccl3 and Ccl5)*, along with UPR-associated genes (e.g *HERPUD*, *DNAJB9* or *SRP54* and *SRP72*), in line with their documented ability to secrete pro-inflammatory molecules^29^ (Fig. 2C). This activation, complemented by the up-regulation of *TSLP* and *IL36*, contributed to the activation of “NF-κB signaling” transcription signature observed in response to SubAB (Fig. 2B, 2D and Supplementary Table S1). In addition to the UPR, gene expression signatures related to “TNF-α via NF-kB signaling” and the “inflammatory response” (Molecular Signatures Database, MSigDB 2020)^33^ were commonly the most statistically relevant pathways up-regulated in the different cell subsets exposed to SubAB. These signatures were particularly prominent in monocytes, B cells, CD4^+^ T cells, CD8^+^ T cells, and NK cells (Fig. 2B and 2D and Supplementary Table S1). This confirms the ability of the toxin to trigger an inflammatory response in PBMC subsets, akin to certain stressors and chemical UPR inducers^10^. Interestingly, the same subset of pro-inflammatory cytokines, was previously described to be enhanced in monocyte derived-DCs upon stimulation with thapsigargin, a potent inhibitor of sarco/endoplasmic reticulum Ca^2+^-ATPases (SERCA), in combination with TLR agonists^34^. The *IL23A* gene was found to be a target of the ER stress-induced transcription factor C/EBP homologous protein (CHOP/DDIT3), which exhibits enhanced binding in the context of both ER-stress and TLR stimulation^34^ and is present in several of the gene signatures induced by SubAB (e.g. mTORC1 signaling) (Supplementary Table S1).

**Figure 2.**
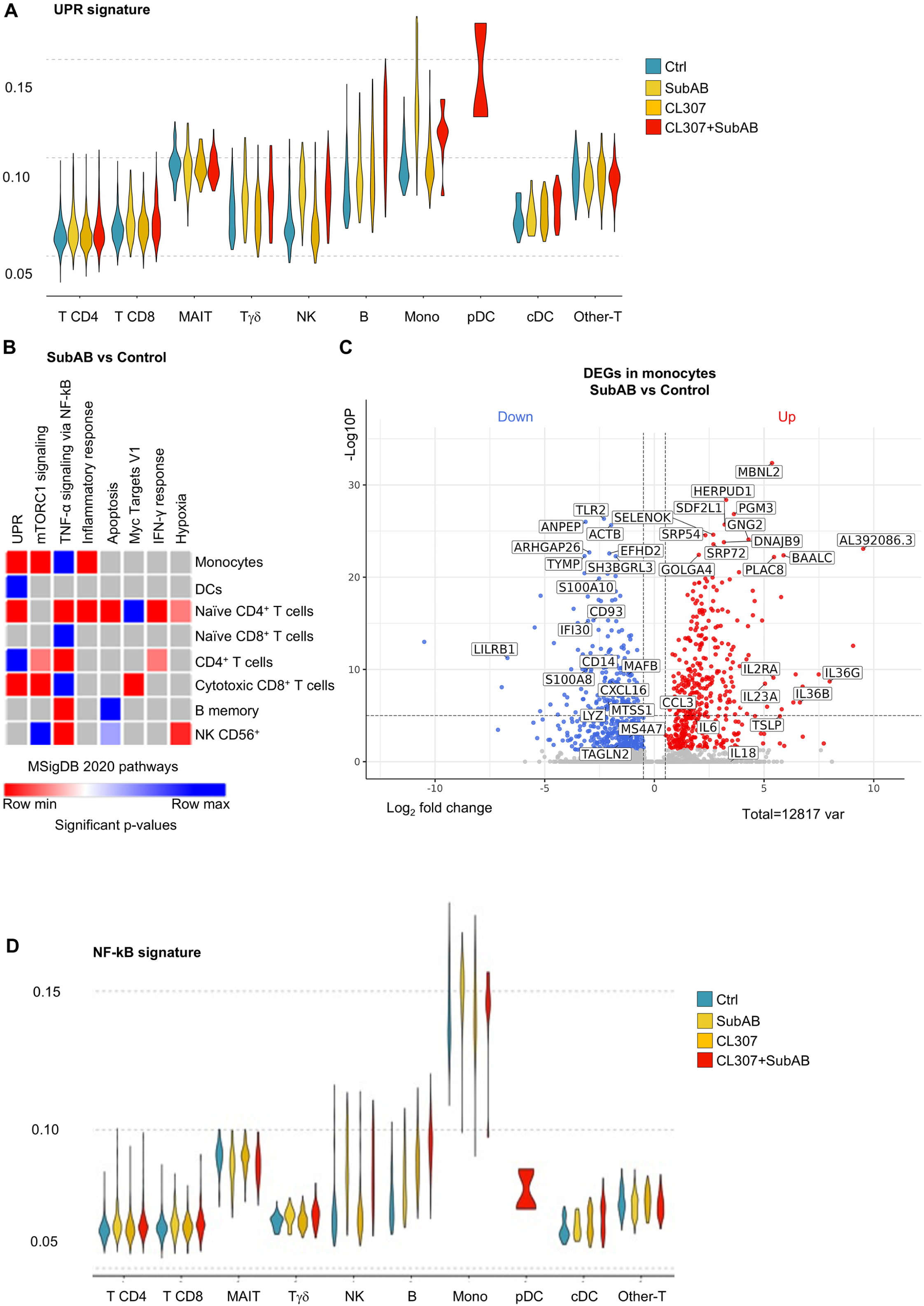
SubAB induces the UPR and a pro-inflammatory response in PBMCs. **(A)** Violin plot of the gene expression signature score of the UPR in the different cell clusters across experimental conditions. **(B)** Heatmap based on P-value significance of the top gene expression pathways (MSigDB 2020) upregulated by SubAB in different PBMC subsets. All significant pathway up-regulations were gradually color coded from lower (blue) to higher (red) significance. When statistical significance is not met, pathways appear in grey. **(C)** Volcano plot representation of the DEGs found in monocytes treated with SubAB compared to control untreated cells. **(D)** Violin plot of the signature score of NF-κB/inflammatory activation in the different cell clusters across experimental conditions.

### scRNA-seq reveals that SubAB exposure influences TLR7 signaling

When scRNA-seq profiles of CL307-treated PBMCs were examined, we observed that TLR7 stimulation induced gene transcription in most cell subsets (Supplementary Fig. S2B), with mostly activation of type-I IFN antiviral responses (Fig. 3A), except for B cells which primarily up-regulated “TNF-α signaling via NF-κB” and the “inflammatory response” (MSigDB 2020)^33^. The DEG response to CL307 observed in most cells involves a combination of direct TLR7 signaling stimulation and activation of the type-I IFN receptor (IFNAR), and signal transducer and activator of transcription 1 (STAT1) signaling after type-I IFN production by unidentified cell subsets. In fact, the IFN-α and IFN-γ pathways both identified using EnrichR show nearly equivalent gene signatures encompassing mostly STAT1-dependent genes and likely representing type-I interferon stimulated genes (ISGs) only (Fig. 3B and Supplementary Table S1). However, one should note that the detection of type-I IFN mRNAs poses a challenge in the context of scRNA-seq analysis of PBMCs and was not detected in any of the datasets examined, preventing the clear identification of IFN-producing cells^30^. Interestingly, co-treatment with SubAB did not affect the IFN-α response, but potentiated the “TNF-α via NF-κB signaling” pathway alongside the UPR in several cell subsets, most notably CD4^+^ T cells. This suggests that the toxin synergizes with the TLR7 signal transduction pathway and can modify downstream transcriptional landscapes (Fig. 3C and Supplementary Table S1). The intensity of this effect was not as acute in monocytes, which induced instead, a pathway called “allograph rejection” corresponding to the induction of some NFKB1 or IL-12-induced gene (*CCND3, CSF2, GADD45B, CD7, BCL, STAT4, CD247, CD3E, CD3D*). Co-treatment with CL307 and SubAB also unveiled a pDC cluster (Fig. 1D and 1E), undetected in other tested conditions (Supplementary Fig. S2A). pDCs, representing only 0.2-0.8% of PBMCs^35^ are nevertheless the principal type-I IFN producing circulating cell subset and their detection by low depth scRNAseq can be challenging. It was striking, however, that the pDC cluster was associated with the highest intensity UPR signature among the different cell subsets analyzed (Fig. 2A). This distinctive signature could potentially mirror a synergy between UPR induction by SubAB and TLR7 stimulation leading to type-I IFN production, as previously suggested for mouse DCs^36^. Overall, SubAB induces the UPR, together with clear pro-inflammatory effects on various human blood cells, but also seems to alter the responsiveness of different subsets, with potentially cross-talks with TLR7 signaling.

**Figure 3.**
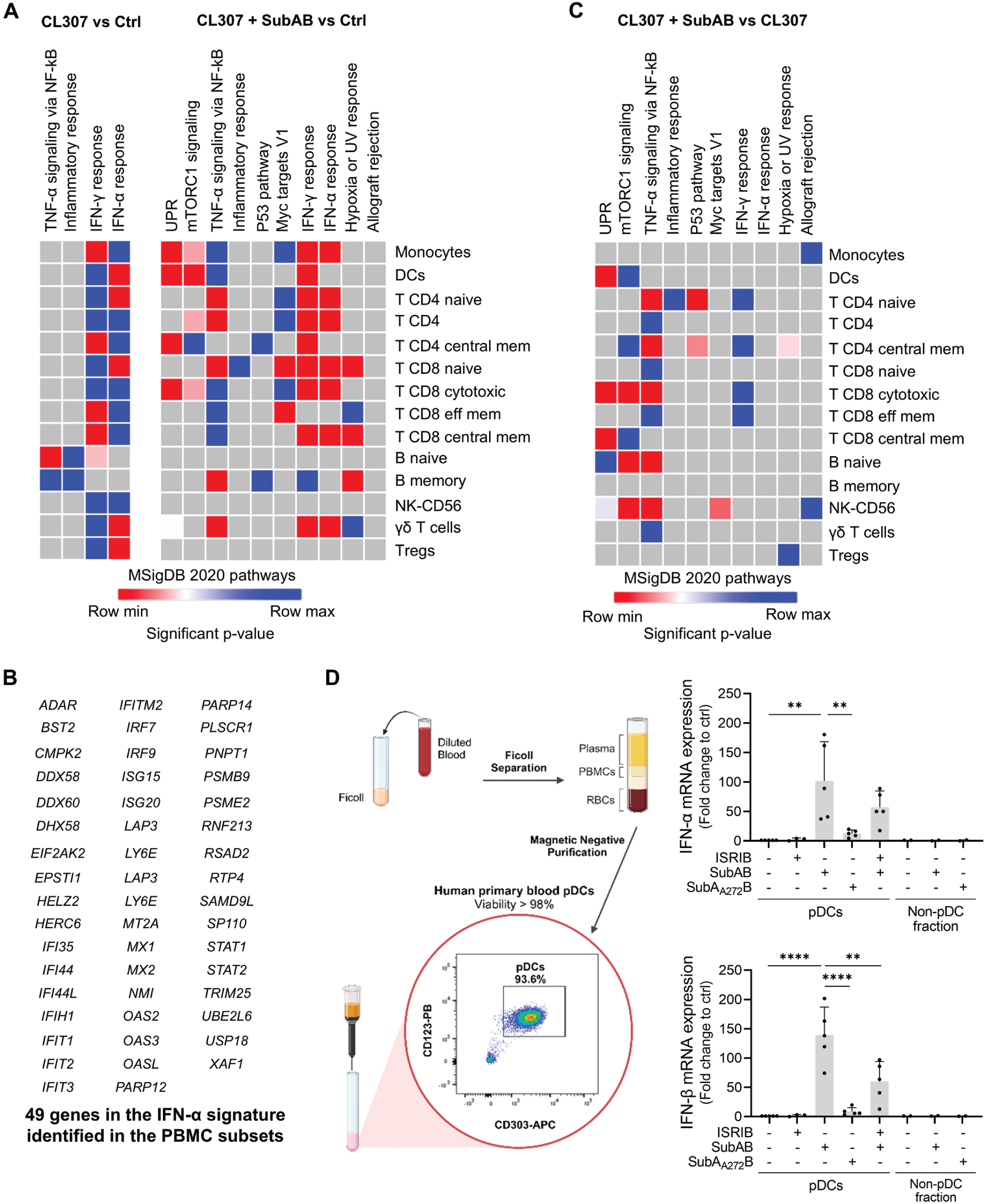
SubAB and CL307 induce pathways leading to both type-I IFN and pro-inflammatory responses. **(A)** Heatmaps calculated on P-value significance of the top gene expression pathways (MSigDB 2020) upregulated by CL307 or SubAB in different PBMC subsets. All significant pathway up-regulations are gradually color coded from lower (blue) to higher (red) significance. When statistical significance is not reached, pathways appear in grey. **(B)** Type-I ISGs up-regulated in CL307-treated PBMCs and being present in both IFN-α and IFN-γ responses signature pathway (MSigDB 2020)**. (C)** Heatmaps based on P-value significance of the top gene expression pathways (MSigDB 2020) upregulated by CL307 and SubAB co-treatment compared to CL307 alone in different PBMC subsets. All significant pathway up-regulations are gradually color coded from lower (blue) to higher (red) significance. When statistical significance is not reached, pathways appear in grey. **(D)** *(left)* Schematic representation of primary human pDCs isolation*. (right)* qPCR monitoring of IFN-α/β mRNA expression of primary human pDCs exposed to 1 µg/ml of SubAB or of proteolytically inactive mutant SubA_A272_B for 6 h alone or combined with the ISR inhibitor ISRIB (750 nM) for 1h. One-way ANOVA followed by Sidak’s multiple comparisons test. Created in BioRender. Pierre, P. (2023) BioRender.com/f62c161.

### SubAB induces type-I IFN in primary blood pDCs

Given the intensity of the UPR observed in pDCs upon treatment with SubAB and CL307, we performed quantitative polymerase chain reaction (qPCR) to measure the activation of type-I IFNs mRNA expression after SubAB treatment in purified human primary blood pDCs. A significant induction of both IFN-α (*IFNA1*) and IFN-β (*IFNAB1*) mRNA expression in response to SubAB alone was observed after 6 h (Fig. 3D). In contrast, the pDC-depleted PBMC fraction did not exhibit any type-I IFN expression (Fig. 3D), confirming that the response was specifically associated with the toxin activity on pDCs. Notably, the proteolytically inactive form of SubAB (SubA_A272_B)^18^ failed to induce type-I IFN expression, underscoring that bacterial toxin uptake is insufficient to elicit a response in primary pDCs and that SubA protease activity is necessary for type-I IFN expression, rather than potential microbe-associated molecular patterns (MAMPs) contaminants. BiP cleavage, and likely the subsequent ISR, are required for type-I IFN induction in this cell type. This is evidenced by the reduced IFN production when co-treated with the small inhibitor of the stress response ISRIB^37^ (Fig. 3D and see scheme Supplementary Fig. S3).

### Human pDCs display active UPR and respond to subtilase cytotoxin

The scarcity of freshly isolated blood pDCs does not allow to perform advanced cell biological and biochemical experiments^38^. Thus, we used the leukemic blastic plasmacytoid dendritic cell neoplasm (BPDCN)-derived cell line CAL-1^39^, which shares many phenotypic and functional properties with freshly isolated human pDCs^33,40,41^. This allowed us to further investigate the potential impact of SubAB on the regulation of type-I IFN expression.

CAL-1 were exposed to increasing concentrations of the PERK inhibitor GSK2656157 (PERKi)^42^ (see scheme Supplementary Fig. S3), and the levels of eIF2α phosphorylation assessed by immunoblot (Fig. 4A). P-eIF2α levels gradually decreased in the presence of increasing doses of the PERKi. Notably, PERKi was optimal at a concentration of 100-300 nM over 6 h, effectively quenching PERK auto-phosphorylation. We generated knock-out CAL-1 cells for the four known *eif2ak* genes (*Hri/EIF2AK1, Pkr/EIF2AK2, Perk/EIF2AK3* and *Gcn2/EIF2AK4*), using CRISPR-Cas9 technology (Supplementary Fig. S4A). Only *Perk* ^-/-^ cells exhibited a marked reduction in eIF2α phosphorylation levels (Supplementary Fig. S4B), confirming the central contribution of PERK for maintaining high levels of eIF2α phosphorylation under steady-state conditions in pDCs^43^. To assess IRE1α activity in CAL-1 cells, we used an IRE1 inhibitor (4µ8C, IRE1i; see scheme Supplementary Fig. S3) and determined levels of XBP1 mRNA splicing by qPCR. Consistent with previous findings in mouse pDCs^8^, CAL-1 cells exhibited a relatively high baseline of XBP1 mRNA splicing which was inhibited by IRE1i treatment (Fig. 4B). Thus, CAL-1 cells, like their primary counterpart, exhibit an ongoing chronic UPR likely initiated during their differentiation^8,9,10,43,44^.

**Figure 4.**
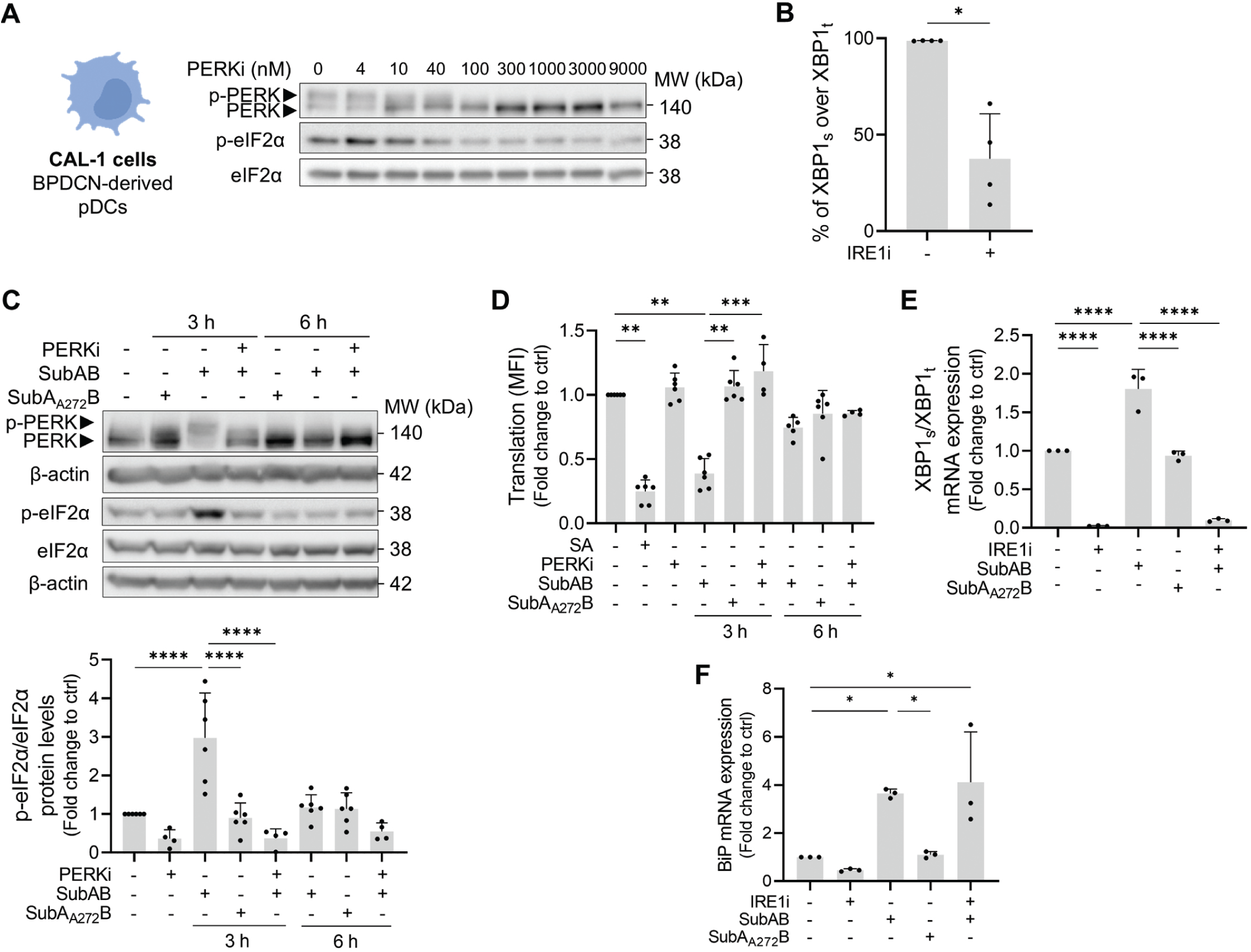
SubAB induces type-I IFN and ER stress in human pDCs. **(A)** Immunoblot detection of p-PERK (upper band), PERK (lower band), p-eIF2α and eIF2α protein levels in CAL-1 cells exposed to increasing concentrations of a PERK inhibitor (PERKi, GSK2656157 for 1 h). **(B)** XBP1 splicing levels in CAL-1 cells treated with the IRE-1α inhibitor (10 μM of IRE1i, 4μ8C, for 1 h) monitored by qPCR. **(C)** *(top)* Immunoblot detection of PERK and eIF2α phosphorylation levels in CAL-1 cells. *(bottom)* P-eIF2α/eIF2α quantification is shown in the bar plot. **(D)** Protein synthesis levels of CAL-1 cells measured by flow cytometry intracellular staining of puromycin incorporation as read-out. Sodium arsenite (500 µM, 30 min) was used as control. **(E)** XBP1_s_/XBP1_t_ and **(F)** BiP mRNA expression monitored by qPCR in CAL-1 cells exposed to SubAB, SubA_A272_B and/or IRE1i. Data was normalized to the housekeeping gene (*Gapdh*). Each dot represents a biological replicate. Data is represented as mean ± SD. (B) Unpaired *t* test with Welch’s correction, (C, E, F) one-way ANOVA followed by Sidak’s multiple comparisons test or (D) Kruskal-Wallis test followed by Dunnett’s multiple comparisons test. Created in BioRender. Pierre, P. (2024) BioRender.com/c99f344.

Next, we evaluated the potential of the SubAB toxin to induce further an ER-stress response in CAL-1 cells. After 3 h incubation with SubAB, a distinctive and complete phosphorylation and activation of PERK, along with subsequent eIF2α phosphorylation, was evident, in contrast to control conditions involving the innocuous SubA_A272_B or PERKi (Fig. 4C). After quantifying the p-eIF2α/eIF2α ratio, it became evident that CAL-1 cells effectively counteracted SubAB’s impact after 6 h stimulation (Fig. 4C). This suggests that, unlike outcomes observed in Vero or HeLa cells^22^, CAL-1 cells possess the ability to rapidly resolve the stress induced by SubAB^45^. This phenomenon recalls the intrinsic resistance of mouse DC subsets to initiate the ISR in response to various chemical ER stressors^43^. By employing the “SUnSET” technique, involving puromycilation followed by flow cytometry detection^46^, we observed a significant inhibition of protein synthesis upon 3 h of SubAB exposure, similar to the effect of sodium arsenite (SA), used as positive control (Fig. 4D). This result is consistent with the increased levels of phosphorylation observed for PERK and eIF2α (Fig. 4C). The impact of the toxin was again strongly attenuated after 6 h of treatment, while PERK inhibition effectively prevented protein synthesis loss (Fig. 4D). SubAB also increased XBP1 mRNA splicing in an IRE1α-dependent manner (Fig. 4E), together with the transcription of the BiP mRNA (Fig. 4F), known to be ATF6-dependent^47^, confirming the broad induction of the UPR mediated by the toxin. Thus, in alignment with primary DCs^43^, CAL-1 cells sustain active protein synthesis despite chronic PERK and IRE1α activities and maintain relative sensitivity to SubAB exposure albeit for a comparatively much shorter duration than other cell lines^22^.

### CAL-1 exposure to SubAB and a TLR7 agonist synergizes to induce IFN-ß

IFN-ß and TNF-α are the two main cytokines produced upon activation of CAL-1 cells^38^. To evaluate the ability of SubAB to induce the expression of these cytokines, we treated cells for 6 h with SubAB alone or in combination with CL307 to recreate the experimental conditions of the PBMC analysis. Upon exposure to SubAB, we observed a 25-fold increase in IFN-ß mRNA expression, which was not observed for SubA_A272_B-treated cells (Fig. 5A). For comparison, stimulation of CAL-1 cells with CL307 resulted in a 100-fold increase of IFN-ß mRNA levels. Interestingly, the combination of SubAB and CL307 exhibited a strong synergistic effect over CL307 alone (≃ x10) on both IFN-ß mRNA expression and protein secretion (Fig. 5A). Accumulation of p-STAT1, a hallmark of CAL-1 activation by CL307 and indicative of IFN-ß production, was distinctly observed upon SubAB exposure alone at 6 h, validating the toxin’s dependent cell activation (Fig. 5B). SubAB did not impact TNF-α mRNA expression (Fig. 5C), suggesting that in pDCs, the toxin triggers specific signaling pathways leading to type-I IFN expression^38^, rather than promoting pro-inflammatory cytokines as observed for some PBMC subsets. To assess the contribution of the different UPR branches to type-I IFN induction, CAL-1 cells were treated with SubAB in the presence or absence of PERKi or IRE1i (Fig. 5D). The treatment with PERKi significantly reduced SubAB-dependent IFN-ß mRNA expression (vs. SubAB alone). IRE1i, in turn, had no significant impact (Fig. 5D), suggesting that PERK activity is the key UPR branch driving induction of IFN-ß mRNA expression in response to SubAB.

**Figure 5.**
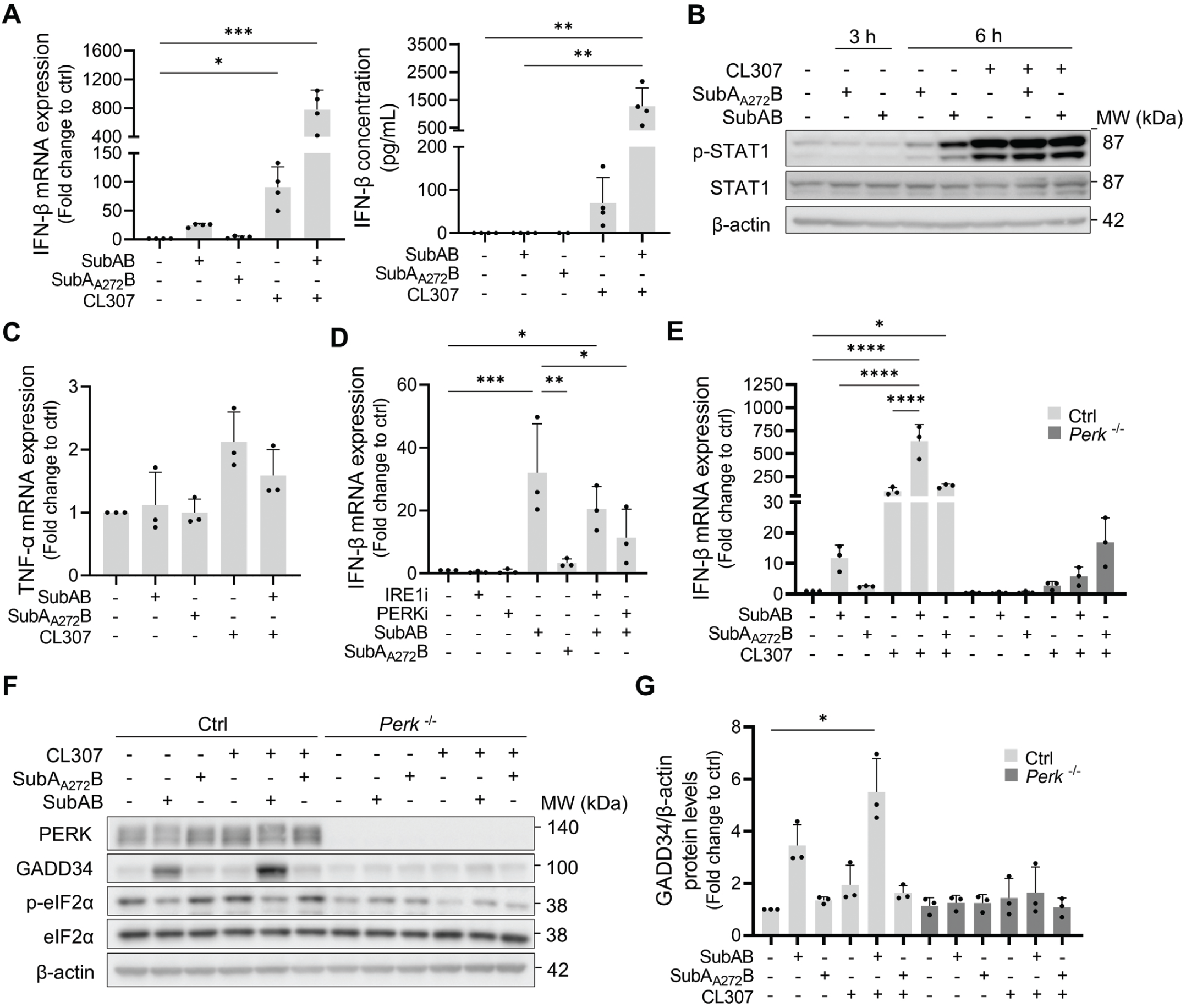
SubAB induces IFN-β expression and synergizes with TLR7 stimulation in CAL-1 cells. CAL-1 cells were treated for 6 h (unless otherwise stated) with 1 µg/ml of SubAB or SubA_A272_B and/or 1μM CL307. **(A)** (*left*) qPCR monitoring of IFN-β mRNA expression and (*right*) IFN-β secretion analyzed by ELISA. **(B)** p-STAT1 levels expression analyzed by immunoblot. **(C)** qPCR monitoring of TNF-α mRNA expression. **(D)** IFN-β mRNA expression was assessed by qPCR in cells pre-treated with PERKi or IRE1i before SubAB exposure. **(E)** IFN-β mRNA expression, **(F)** PERK, GADD34 and p-eIF2α protein levels and **(G)** quantification of GADD34 protein levels in control (Ctrl) and *Perk*^-/-^ CAL-1 cells. Immunoblots are representative of at least 3 independent experiments. Data is represented as mean ± SD and each dot represents a biological replicate. (A, C, G) Kruskal-Wallis test followed by Dunnett’s multiple comparisons test or (D, E) one-way ANOVA followed by Sidak’s multiple comparisons test.

To confirm the involvement of PERK in SubAB-induced IFN-ß mRNA expression, we compared the effect of SubAB on control or *Perk*^-/-^ CAL-1 cells (Fig. 5E). Both SubAB and CL307 triggered IFN-ß mRNA induction and exhibited synergistic effects in control cells, but the effect of SubAB was abolished by PERK inactivation (Fig. 5E). *Perk*^-/-^ cells were not able to initiate an ISR in response to the toxin, as indicated by low p-eIF2α levels and the lack of expression of GADD34/PPP1R15a, a phosphatase-1 adaptor known to target eIF2α phosphorylation and to depend on the ISR for its synthesis^48^ (Fig. 5F-G). The induction of GADD34 after 6 h of SubAB exposure likely accounted for the dephosphorylation of eIF2-α and the restoration of protein synthesis (Fig. 4C-D, and 5F-G), facilitating the substantial secretion of IFN-ß in cells co-treated with SubAB and CL307^36^. Moreover, PERK inactivation also impaired the ability of CAL-1 cells to respond to CL307, as evidenced by a reduction in IFN-ß expression in *Perk*^-/-^ cells (Fig. 5E). Thus, the activity of PERK is key for driving IFN-ß expression in CAL-1 cells in response to both SubAB and CL307, albeit with different efficiency and probably via distinct mechanisms.

### BiP inhibition by HA15 drives IFN-ß mRNA transcription in CAL-1 cells

The increase of IFN-ß mRNA following SubAB exposure might stem from the stabilization of mRNA within Ras GTPase-activating protein-binding protein 1 (G3BP1)- containing stress granules (SGs)^49,50^, rather than just an elevation in transcription levels. In line with increased p-eIF2α levels (Fig. 4C), SubAB exposure drove the assembly of G3BP1-containing SGs (Fig. 6A). However, concomitant transcription inhibition using actinomycin D prevented IFN-ß mRNA up-regulation and even promoted decay over time (Fig. 6B). This indicates that active SubAB triggers *de novo* IFN-ß transcription through a complex biochemical response including PERK-dependent eIF2α phosphorylation, rather than mRNA stabilization and storage in SGs. We next explored whether selective inactivation of BiP could account for SubAB’s efficacy. We used HA15, a thiazolidinedione compound recognized for its exclusive targeting of BiP and induction of ER stress^51^. Like with SubAB, CAL-1 exposure to incremental concentrations of HA15 for 6 h, led to eIF2α phosphorylation and GADD34 synthesis (Fig. 6C), accompanied with an increase in type-I IFN mRNA and p-STAT1 levels, albeit with lesser intensity for the later (Fig. 6D-E). Thus, targeting of BiP stands as a pivotal driver in orchestrating this process, whether instigated by the toxin or through chemical agents, like HA15.

**Figure 6.**
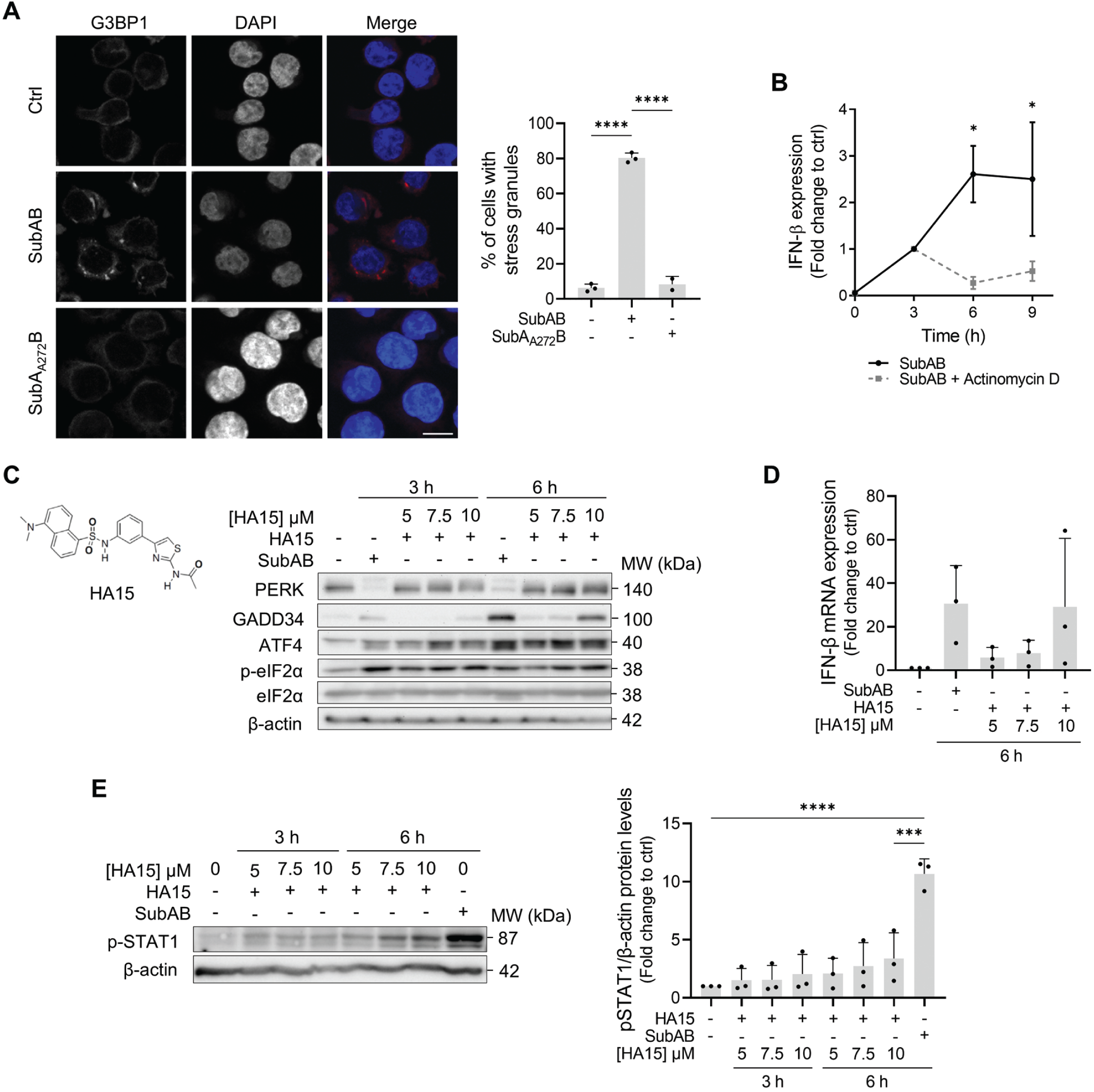
Inactivation of BiP drives *de novo* synthesis of IFN-β. **(A)** (*left*) Confocal immunofluorescence microscopy for G3BP1 (red) and nuclei (DAPI, blue) in CAL-1 cells treated with SubAB for 6 h. (*right*) The percentage of cells with SGs is presented in the graph. At least 100 cells were analysed per condition. Scale bar 10 µm. **(B)** IFN-β mRNA expression analysed by qPCR through time in CAL-1 cells exposed to SubAB and actinomycin D (5 µg/mL, added 3 h after SubAB). **(C-E)** CAL-1 cells were treated with increasing concentrations of the small compound HA15 for the indicated timepoints. **(C)** PERK, GADD34, ATF4 and p-eIF2α protein levels were analysed by immunoblot, **(D)** IFN-β mRNA expression by qPCR and **(E)** STAT1 phosphorylation by immunoblot. Immunoblots are representative of 3 independent experiments. Data is represented as mean ± SD and each dot represents a biological replicate. (A, C, G) Kruskal-Wallis test followed by Dunnett’s multiple comparisons test or (D, E) one-way ANOVA followed by Sidak’s multiple comparisons test.

### Induction of IFN-ß transcription by SubAB is STING and p-eIF2α dependent

Different viral sensing networks, such as the cGAS/STING and RIG-I/MDA5-like RNA (RLR)/mitochondrial antiviral signaling protein (MAVS) signaling pathways, have been identified as key initiators of type-I IFN expression^52^. These networks rely on the adaptor proteins STING (TMEM173) and MAVS (also known as VISA, IPS-1, Cardif) to receive antiviral signals from distinct cytosolic RNA- (RLRs) or DNA-sensors (cGAS)^53^. This activation subsequently prompts the TBK1/IKKε/IRF3 signaling cascade (see scheme Fig. 7A)^54^. Regardless of the origin of the signaling stimulus, the TBK1/IKKε axis holds a pivotal role in facilitating IRF3 phosphorylation, its translocation into the nucleus, and the transcription of IFN-ß^55^. To dissect the influence of SubAB on TBK1/IKKε signaling and its impact on IFN-ß mRNA expression, we employed the TBK1 inhibitor MRT67307 (TBK1i; see scheme Fig. 7A)^56^. TBK1i treatment efficiently suppressed SubAB-triggered IFN-ß expression (Fig. 7B), underlining the central involvement of TBK1, and suggesting that IFN-ß mRNA induction likely stems from either STING or MAVS activation by SubAB.

**Figure 7.**
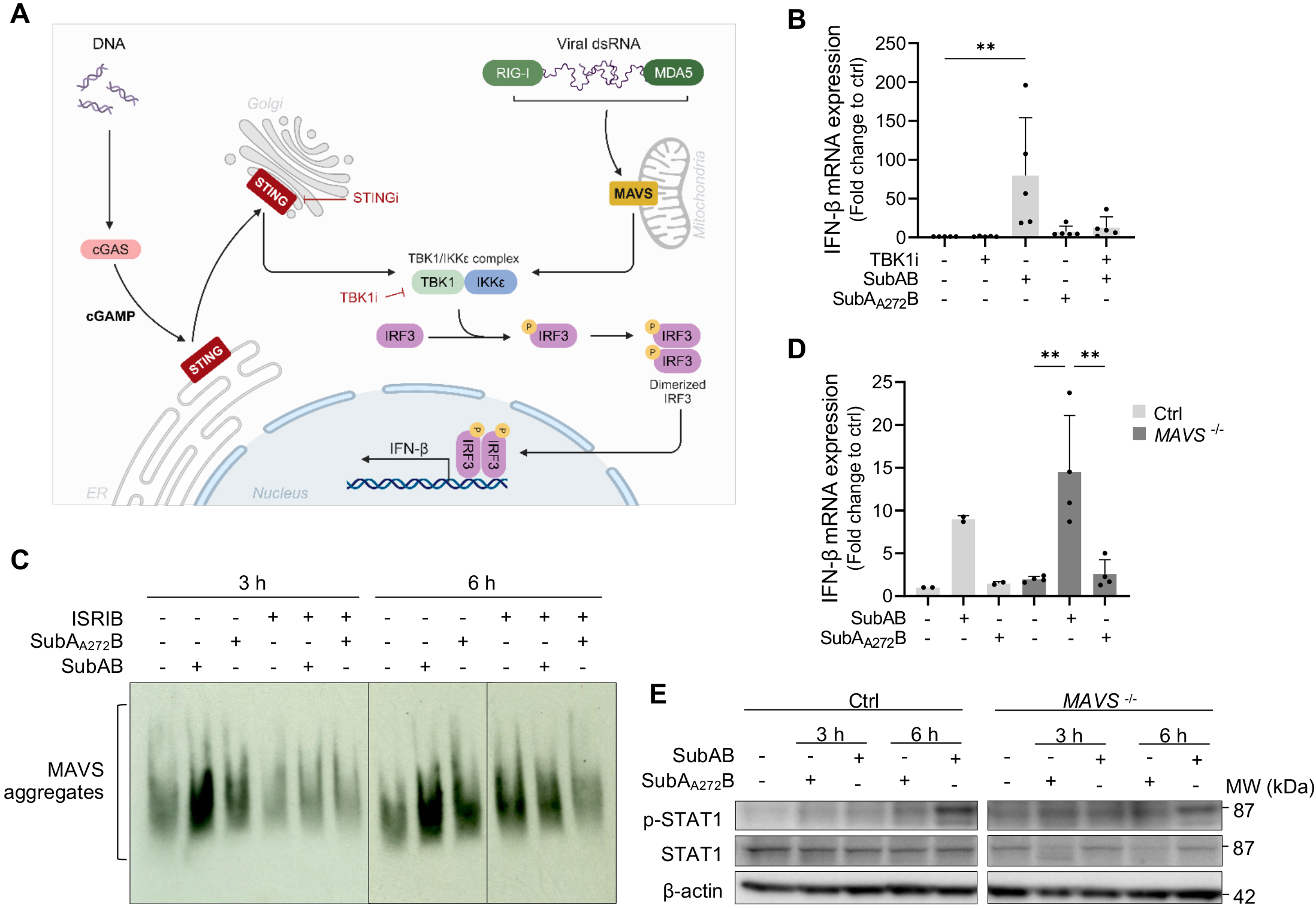
SubAB drives IFN-β expression independently of MAVS signaling. **(A)** Schematic representation of the cGAS/STING and RIG-I/MDA-5/MAVS (RLR) pathways. **(B)** IFN-β mRNA expression in CAL-1 cells pre-treated with TBK1i (2 µM) for 1 h and exposed to SubAB for 6 h. **(C)** MAVS aggregation was evaluated by SDD-AGE in cells exposed to ISRIB and/or SubAB. Representative image of n=5 biological replicates. **(D, E)** Control (Ctrl) or *MAVS*^-/-^ CAL-1 cells were exposed to SubAB. **(D)** IFN-β mRNA expression at 6 h and **(E)** p-STAT1 protein levels were analyzed. Data is represented as mean ± SD and each dot represents a biological replicate. (B) Kruskal-Wallis test followed by Dunnett’s multiple comparisons test or (D) ordinary one-way ANOVA followed by Sidak’s multiple comparisons test. Created in BioRender. Pierre, P. (2024) BioRender.com/h19l873.

Upon the detection of nucleic material by the RLRs, MAVS undergoes the development of a prion-like structure with functional properties^57^. This structure acts as a central hub, facilitating the assembly of a signalosome [signaling MAVS organizing centers (SMOC)], which subsequently triggers the activation of TBK1 and IKKε (Fig. 7A). It has been noted that the assembly and signaling of SMOCs are reliant on eIF2α phosphorylation and ATF4^58^. Treatment with SubAB for 3 and 6 h led to a notable accumulation of aggregated MAVS and potentially SMOC assembly, as evidenced through semi-denaturing detergent agarose gel electrophoresis (SDD-AGE) (Fig. 7C). MAVS aggregation was found to be sensitive to ISRIB, reinforcing the significance of the ISR and eIF2α phosphorylation in this context. However, SubAB retained the capability to induce type-I IFN expression (Fig. 7D) and drive STAT-1 phosphorylation (Fig. 7E) in *MAVS*^-/-^ cells (Supplementary Fig. S5A), while these cells were, as expected, unresponsive to cytosolic dsRNA (poly I:C) delivery (Supplementary Fig. S5B). MAVS is therefore not required to drive IFN-ß expression in response to SubAB. Moreover, the levels of MAVS aggregation observed in response to the toxin are insufficient to activate TBK1, contrasting with previous reports based on the ectopic expression of MAVS mutants^59,60^.

TBK1 activation may also be orchestrated by STING, an ER-resident signaling adaptor subject to stringent regulation of expression and trafficking (see scheme Fig. 7A)^61^. This intricate control aims to curtail auto-activation and excessive IFN-ß production, which could potentially trigger interferonopathies^62,63^. To investigate the potential involvement of the STING pathway on SubAB-induced IFN expression, we used the STING inhibitor H-151 (STINGi, see scheme Fig. 7A)^63^ in combination with SubAB. Up-regulation of IFN-ß mRNA was reduced upon STING inhibition (Fig. 8A). As anticipated by the moderate levels of IFN expression, SubAB only mildly altered STING’s homeostasis after 3 h (Fig. 8B), suggesting limited Golgi translocation and endosomal degradation compared to the activation levels observed after 3 h of cytosolic cGAMP exposure used as a positive control (Supplementary Fig. S5C)^53^. To confirm our observation, *STING1*^-/-^ CAL-1 cells were generated (Supplementary Fig. S5C-D), and exposed to SubAB. Complete STING inactivation impaired IFN-ß production (Fig. 8C) and STAT1 phosphorylation (Fig. 8D) upon SubAB uptake, while it did not markedly reduce CAL-1’s ability to synergize with TLR7 stimulation (Fig. 8E). This suggests that STING is required for the response to SubAB alone but not for the synergy observed with TLR7 signaling, although both pathways appear to rely on PERK activation to drive increased type-I IFN expression. Surprisingly, bypassing eIF2α phosphorylation using ISRIB did not affect the capacity of SubAB to synergize with CL307 (Fig. 8F). This further suggests that this event depends on an alternative function of PERK, rather than protein synthesis down-modulation, which seems key only to control the STING signaling.

**Figure 8.**
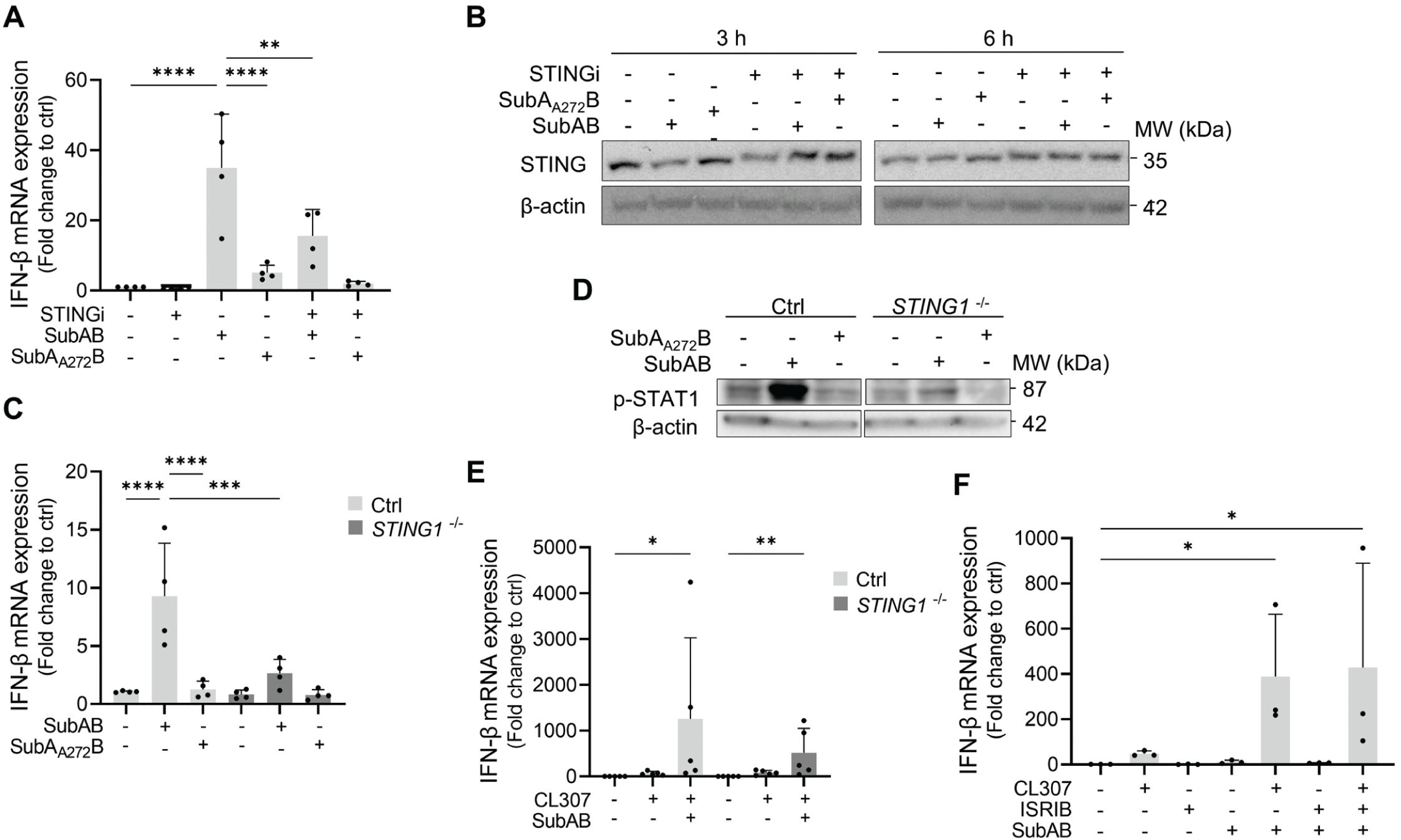
STING inactivation impairs IFN-β expression triggered by SubAB. (A, **B)** CAL-1 cells were pre-treated for 1 h with the STING inhibitor H-151 (STINGi; 0.5 µM) before exposure to SubAB for 3 or 6 h. **(A)** IFN-β mRNA expression (6 h), **(B)** STING protein level in Control (Ctrl) CAL-1 cells during indicated time. Immunoblot representative of 2 biological replicates. **(C, E, F)** IFN-β mRNA expression and **(D)** P-STAT1 levels in Control (Ctrl) and *STING1*^-/-^ CAL-1 cells exposed to SubAB alone **(C, D)** or in combination with CL307 (**E)** or ISRIB **(F)**. Immunoblot representative of 3 biological replicates. Data is represented as mean ± SD and each dot represents a biological replicate. Ordinary one-way ANOVA followed by Sidak’s multiple comparisons test, except for (**E**), where Kruskal-Wallis test followed by Dunnett’s multiple comparisons test was performed.

### The ubiquitin-binding adapter TAX1BP1 is down-regulated in response to SubAB

We hypothesized that SubAB might influence not only translation regulation and STING activation, but also various pathways associated with innate sensing. Previous research has demonstrated that Unc-51 like autophagy activating kinase 1 (ULK1) can phosphorylate and inhibit STING, resulting in the suppression of IRF3 activation^64^. However, the inactivation of ULK1 in CAL-1 cells did not result in any discernible alteration of IFN-ß induction (Supplementary Fig. S6A-B). This suggests that the ULK1 pathway, typically associated with nutrient starvation and autophagy, is not triggered during the SubAB-dependent UPR. To explore whether SubAB affected macroautophagic flux and known autophagy receptors, we monitored autophagy using microtubule-associated proteins 1A/1B light chain 3 (ATG8/LC3) flux over time in the presence of chloroquine (CQ) or bafilomycin A (Baf A)^65^ (Supplementary Fig. S6C). SubAB did not impact autophagy, while conversely, inactivation of *ATG5* in CAL-1 cells (Supplementary Fig. S6D)^65^ had no influence on IFN-ß induction nor STAT1 phosphorylation by SubAB (Supplementary Fig. S6E-F). This further underscores that macroautophagy does not participate in this process, not even indirectly, by, for example, preventing the clearance of activated STING^15^.

Considering the pivotal role of various selective ubiquitin-binding sequestosome-like autophagy receptors (SLRs) like protein sequestosome 1 (p62/SQSTM1), neighbor of *Brca1* gene (NBR1), nuclear dot protein 52 (NDP52), optineurin, or TAX1BP1 in the regulation of various innate immune signaling pathways^66,67,68^, we investigated the impact of SubAB treatment on their expression through immunoblot analysis (Fig. 9A). Most of the SLRs tested remained relatively unaffected by the toxin with the exception of p62 and TAX1BP1, which experienced down-modulation after 3 h of SubAB exposure (Fig. 9A and Supplementary Fig. S7A). TAX1BP1 displayed however the most acute and enduring loss (6 h) despite slightly augmented mRNA levels over time (Fig. 9B). TAX1BP1 disappearance could be entirely prevented in *Perk*^-/-^ cells or upon ISRIB treatment (Fig. 9C-D), but was not influenced by STING deletion (Fig. 9E), effectively disconnecting STING activation from the ISR, which was uniquely responsible for triggering TAX1BP1 disappearance.

**Figure 9.**
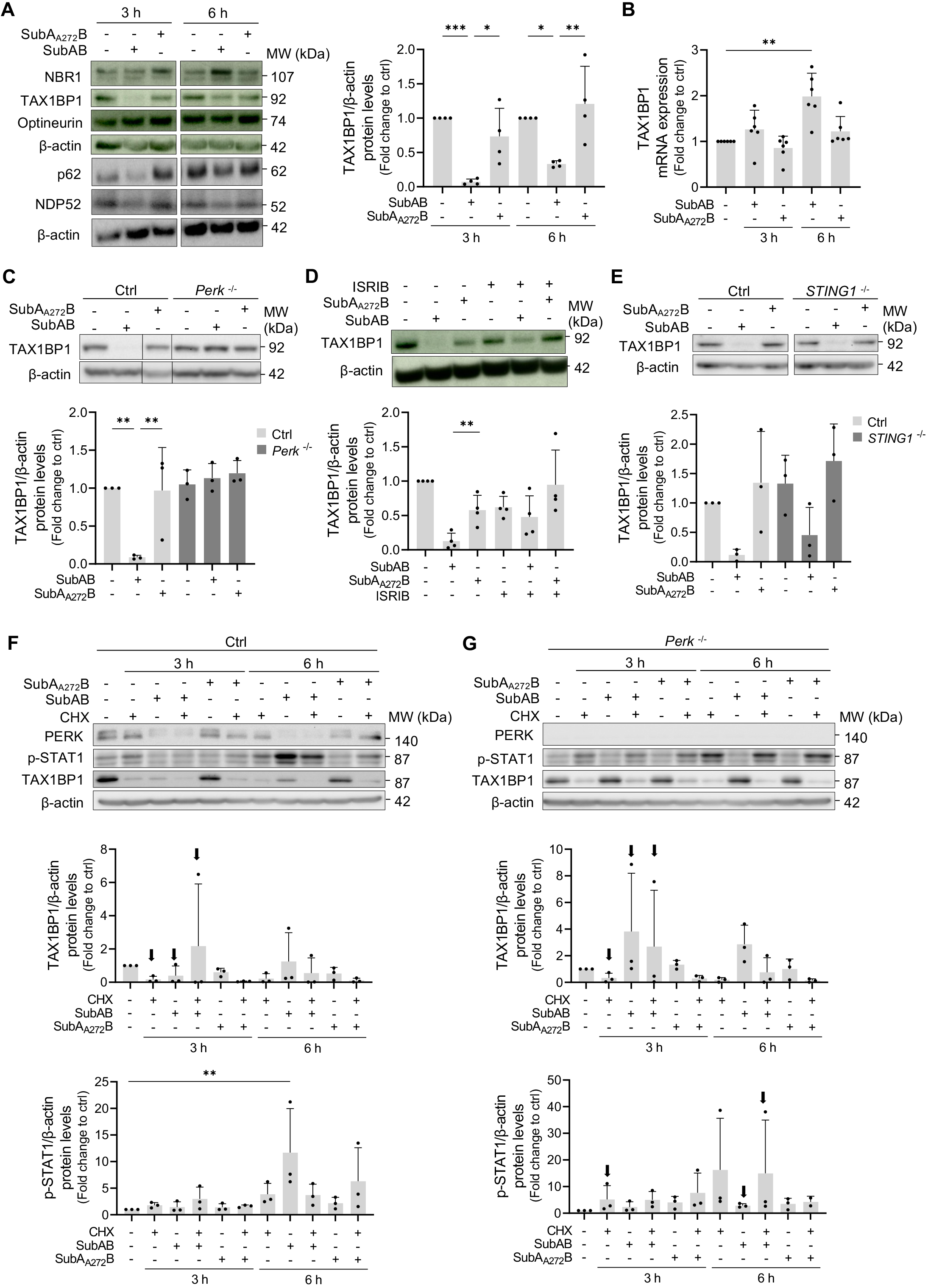
Translation inhibition-dependent loss of TAX1BP1 upon SubAB treatment. **(A)** (*left*) CAL-1 cells were treated with SubAB or inactive toxin for the indicated timepoints, and the protein levels of SLRs were evaluated by immunoblot as indicated. (*right*) Quantification for TAX1BP1 levels is shown. **(B)** TAX1BP1 mRNA levels were analyzed by qPCR. **(C-E)** Immunoblot monitoring of TAX1BP1 protein levels in **(C)** Control (Ctrl) and *Perk*^-/-^ cells, **(D)** in combination with ISRIB treatment and **(E)** Ctrl and *STING1*^-/-^ CAL-1 cells exposed to SubAB for 3 h. Quantification is shown below the representative blots, n=3. **(F-G)** Immunoblot monitoring of PERK, p-STAT1 and TAX1BP1 protein levels in **(F)** Ctrl and **(G)** *Perk*^-/-^ CAL-1 cells exposed to cycloheximide (CHX) for the indicated timepoints. Quantification of p-STAT1 and TAX1BP1 levels is shown below representative blots, n=3. Arrow point at key conditions. Data is represented as mean ± SD and each dot represents a biological replicate. (A, C-E) Ordinary one-way ANOVA followed by Sidak’s multiple comparisons test or (B, G, I) Kruskal-Wallis test followed by Dunnett’s multiple comparisons test.

We explored the relationship between the ISR and TAX1BP1 levels by investigating the impact of inhibiting protein synthesis using cycloheximide (CHX) in both control and *Perk* ^-/-^ cells. CHX treatment led to the rapid loss of TAX1BP1 in both cell types (Fig. 9F and 9G). An inverse correlation between p-STAT1 levels and TAX1BP1 expression was also observed (Fig. 9F and 9G). Importantly, the inhibition of protein synthesis alone could induce STAT1 phosphorylation, particularly in *Perk*^-/-^ cells, suggesting an important role of protein synthesis in negatively controlling the IFN expression and signaling at steady state and upon SubAB exposure.

### Selective regulation of negative feedback regulators by SubAB

Among SLRs, TAX1BP1 has been shown to promote MAVS aggregates clearance^69,70^ and importantly, to play a key role in the negative regulation of NF-kB and of TBK1/IRF3 signaling by acting in concert with the ubiquitin-editing enzyme A20/TNFAIP3 (see scheme Supplementary Fig. S7B and Supplementary Table S2)^71,72,73^. We tested the effect of SubAB treatment on the expression of other known negative feedback regulators of inflammatory signaling^74^ by qPCR and immunoblot (Fig. 10A-B, Supplementary Fig. S8A). Out of the 12 regulators tested, mRNA expression of *IFIT1, SOCS1* and *DUSP1* was strongly up-regulated by SubAB (Fig. 10A), with some others, like *CYLD*, *NFKBIA//IKB-α, SOCS3 or A20/TNFAIP3* reflected the signaling synergy between SubAB and CL307. At the protein level only IFIT1 was found to be up-regulated significantly upon co-exposure to SubAB and CL307, the rest of the regulators tested remaining relatively stable irrespective of the intensity of their mRNA expression (Fig. 10B and Supplementary Fig. S8A). PIAS3, which is known to inhibit STAT3, had the tendency to be downmodulated by SubAB, although with no statistic relevance across experimental repeats. Thus, among the negative feedback regulators of inflammatory and IFN signaling tested in CAL-1 cells, only TAX1BP1 significantly required de novo synthesis to maintain its active levels during SubAB-induced ISR. By inhibiting the proteasome, using MG132, and endocytosis or autophagy degradation using VPS34-IN1, a potent inhibitor of class III PtdIns(3)P-kinase VPS34^75^, we could show that TAX1BP1 loss was prevented by VPS34-IN1 treatment but not by MG132 (Supplementary Fig. S8B-C), pointing at a role for endosomal degradation and most likely endosomal microautophagy for its rapid turnover^76^. Thus, TAX1BP1 expression could be constantly dampening STING signaling and its fast disappearance upon SubAB exposure could contribute to enhance type-I IFN expression in steady state and SubAB-exposed pDCs (see scheme Supplementary Fig. S7B).

**Figure 10.**
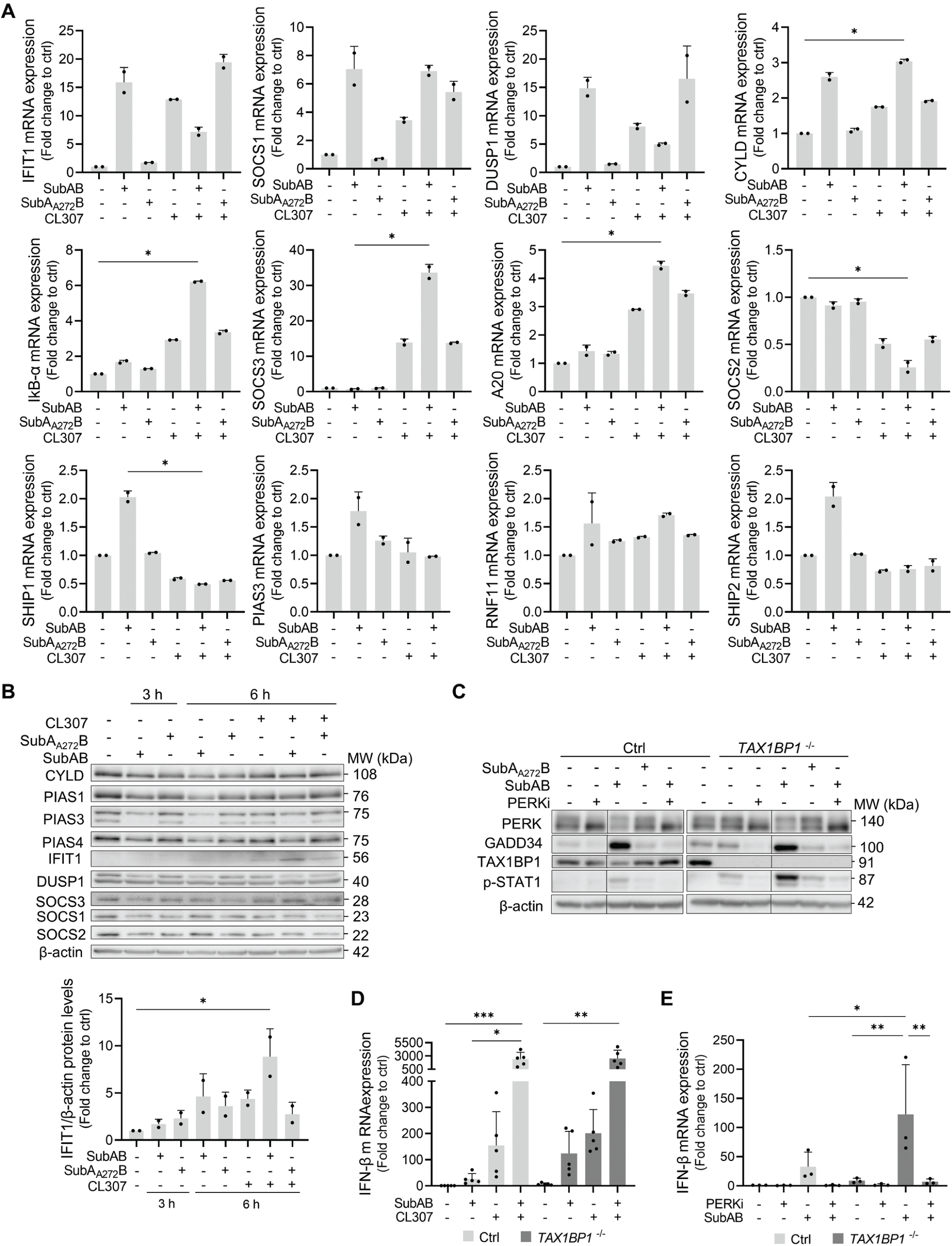
SubAB impacts inflammatory signaling negative feedback regulators. Expression levels of selected inhibitors of innate and cytokine receptors signaling were monitored in cells treated with SubAB and/or CL307 for the indicated timepoints. **(A)** mRNA expression (6 h) by qPCR and **(B)** protein levels by immunoblot and quantification of IFIT1 expression. **(C)** Immunoblot monitoring of PERK, GADD34, TAX1BP1 and p-STAT1 protein levels in Control (Ctrl) and *TAX1BP1*^-/-^ CAL-1 cells exposed to SubAB (3 h) in absence or presence of pre-treatment with PERKi. **(D)** Induction of IFN-ß mRNA expression monitored by qPCR in *TAX1BP1*^-/-^ CAL1 in response to SubAB, CL307 or both (6 h). **(E)** Induction of IFN-ß mRNA expression monitored by qPCR in *Tax1bp1*^-/-^ CAL1 in response to SubAB (6 h) in absence or presence of PERKi. Data is represented as mean ± SD and each dot represents a biological replicate. Immunoblot representative of (B) 2 or (C) 3 biological replicates. (A-D) Kruskal-Wallis test followed by Dunnett’s multiple comparisons test or (E) one-way ANOVA followed by Sidak’s multiple comparisons test.

### TAX1BP1 loss promotes STING activation in response to SubAB

We next tested whether TAX1BP1 protein loss could promote type-I IFN expression upon SubAB or CL307 exposure by generating *TAX1BP1*^-/-^ CAL-1 cells (Supplementary Fig. S8D). Compared to steady state control, *TAX1BP1*^-/-^ cells displayed limited levels of STAT1 phosphorylation in absence of stimulation (Fig. 10C), probably reflecting some minimal type-I IFN expression (Fig. 10D-E). *TAX1BP1*^-/-^ CAL-1 were however hypersensitive to SubAB stimulation by inducing robust IFN-ß mRNA expression and STAT1 phosphorylation (Fig. 10C-D). Despite this clear enhancement, CL307 signaling, or the synergy observed with SubAB upon co-stimulation, were not affected by *TAX1BP1* deletion, reflecting our previous observations (Fig. 10D). Moreover, pharmacological inhibition of PERK suppressed the enhancement of IFN-ß and p-STAT1 at steady state and upon SubAB treatment in *TAX1BP1*^-/-^ CAL-1 cells (Fig. 10C and 10E), suggesting that, in addition of decreasing TAX1BP1 protein levels, additional PERK activity is clearly required to promote STING and TLR7 signaling.

In summary, our findings reveal three parallel but complementary pathways induced by BiP cleavage in pDCs. The first pathway moderately activates STING, likely within the ER, resulting in limited downstream activation of TBK1 and IRF3, which is normally kept under control by negative regulators like TAX1BP1. The second pathway activates PERK, which inhibits cap-mediated translation and impacts TAX1BP1 expression and potentially other unidentified signaling molecules, strongly amplifying STING-dependent TBK1 and IRF3 activation and culminating in robust type-I IFN production in response to SubAB. The third pathway operates independently of STING activation and TAX1BP1 loss, with PERK impacting other unidentified regulatory node(s) that amplify TLR7 signaling upon CL307 stimulation, regardless of eIF2α phosphorylation and protein synthesis inhibition.

## DISCUSSION

SubAB appears to be one of the few identified physiological compounds capable of triggering ER stress. It proves to be a valuable tool to investigate the role of BiP and the UPR in a variety of cellular functions. A direct link between the UPR induction and type-I IFN expression has been, until now, tedious to demonstrate, since most pharmacological ER stressors alone do not trigger type-I IFN production, which was mostly observed in synergy with the activation of *bona fide* innate microbe sensing pathways in different cell types^12^. Importantly, SubAB is lethal for mice and induces pathological features seen during HUS^77^. SubAB’s impact on immune cells could contribute to STEC infection pathogenesis, although the actual role of the toxin in human disease pathogenesis had to be established. We took advantage of the development of scRNA-seq technologies to profile PBMCs from healthy donors exposed to SubAB. Our results show that SubAB triggers the UPR in different cell subsets, the extent and the intensity of which are cell type specific and have profound differences in their outcome. The ISR induced by SubAB polarizes the inflammatory response in toxin-exposed monocytes, while inducing the preferential loss of the CD16^+^ cells. This effect could explain some immunoregulatory aspects of STEC infection via enhanced IL-23 and thymic stromal lymphopoietin (TSLP) expression, and loss of non-classical monocytes capable of promoting neutrophils adhesion at endothelial interfaces^78^. Moreover, we uncovered that DCs display unusual features, allowing them to resist the toxin effect and reestablish protein synthesis and homeostasis quite rapidly, similarly to what has been observed in different mouse DC subsets^43^, but differently from Vero cells^22^.

SubAB is able to induce type-I IFN expression in pDCs, alone or in strong synergy with TLR7 activation. Our work demonstrates that SubAB does not only activate a PERK-dependent ISR after BiP cleavage, but also stimulates the STING pathway, via a yet undefined mechanism. Concomitant activation of these two parallel pathways is necessary for an efficient induction of type-I IFN production. Given that the HA15 compound, that also targets BiP^51^, partially phenocopies SubAB effects, BiP/HSPA5 availability appears to play a central role to prevent basal STING activation directly or indirectly in steady state pDCs. Interestingly, among few other ER resident proteins, ATF6, one of the three UPR sensors that also binds BiP and controls its expression, was recently shown to interact with STING in cGAMP-stimulated U937 cells^79^, suggesting that BiP could have additional regulatory functions than contributing to protein folding and serving as a sensor to trigger the UPR.

Given that professional type-I IFN producing pDCs are characterized by an extremely developed ER network and, like for plasmocytes, depend partially on XBP1 for their differentiation^10^, they might have acquired distinct features, which renders them resistant to SubAB and allow for significant type-I IFN production in response to BiP cleavage. Nevertheless, the transient protein synthesis reduction caused by PERK-dependent eIF2α phosphorylation, functions as an amplification signal required to allow STING activation, while PERK activation can synergize with TLR7 signalling to favour type-I IFN expression by pDCs. These observations may be extrapolated to other types of TLR activation in various cell types^80^, explaining the significative induction of pro-inflammatory pathways in several PBMC subsets upon SubAB exposure.

Translation arrest can be considered as a sign of dysbiosis, which when *bona fide* microbe sensors are activated, leads to signal potentiation^81^. We, and others, described and theorized this phenomenon during double-stranded RNA-dependent protein kinase (PKR/eIFK2) sensing of viral infection and IFN production through augmented IRF3 and NF-kB activation^82,83,84,85^. Importantly, the fact that pDCs cope with SubAB induced ER-stress and rapidly restore protein synthesis levels after PERK activation and upon TLR7 stimulation, allows the efficient translation of the type-I IFN mRNAs whose accumulation was exacerbated in the first hours of ISR induction. Supporting this idea, previous research in mice have proposed that TLR4 activation in macrophages undergoing an ISR, suppresses CHOP induction and protein synthesis inhibition, preventing apoptosis in activated cells ^80^. Moreover, ATF4 binds interferon regulatory factor 7 (IRF7) and prevents type-I IFN transcription^86^. Thus, for pDCs, limiting the consequences of the UPR during time, might be required to optimize type-I IFN transcription as well^43^.

Differently from PKR activation, SubAB-dependent PERK activation affects only moderately the protein levels of the non-exhaustive list of 15 negative feedback regulators tested, at the exception of TAX1BP1, which is rapidly lost (3 h). Although BiP disruption alone is not sufficient to trigger efficiently productive STING activation, we showed that the concomitant loss of the negative feedback exerted by the TAX1BP1/A20 complex^73^ (see scheme Supplementary Fig. S7B), results in an amplification of STING signalling and IFN production. However, in absence of TAX1BP1, PERK activation and potentially the loss of other unidentified negative regulators, are still required to fully achieve IFN expression, as well as trigger synergy with TLR7. We showed that SubAB promotes MAVS aggregation, which is in line with TAX1BP1 controlling its clearance and with the importance of translational regulation for MAVS biochemistry^59,60^. However, differently from previous reports using ectopic expression experiments^59,60^, these changes were insufficient to support auto-activation and type-I IFN production in stressed cells. Although one would expect a strong synergy with RLR stimulation, if these receptors were stimulated concomitantly to BiP’s disruption by pharmacological or viral compounds.

We therefore provide evidence that SubAB promotes cytokine production in key immune cells. We also show how the ISR and associated translation inhibition can synergize with innate sensor activation to bypass negative feedback pathways and amplify cytokine production and probably other immunostimulatory features of immune cells like DCs or B cells. STING appears to be the main driver of IFN-ß mRNA transcription in response to SubAB, supporting the view that STING and PERK are engaged in a crosstalk necessary for their respective activation^17^. These results have also implications on how interferonopathies could be caused by pDC and B cell exposure to different acute ER-stress inducers like bacterial toxins^87^ or metabolic insults in susceptible individuals. Supporting this hypothesis, a mutation in the *TAX1BP1* gene was recently found in a pedigree of a patient family affected by systemic lupus erythematosus (SLE)^27^. The TAX1BP1 protein is expressed at low levels in the PBMCs of SLE patients and shows a negative correlation with disease activity and autoantibody levels. Interestingly silencing of *TAX1BP1* in THP-1 monocytes led to an up-regulation of CD14 and loss of CD16 expression^27^, echoing our results on SubAB-treated monocytes. We anticipate that in families susceptible to immune deficiencies, variants for genes involved in the UPR will also be characterized in the near future^30^.

## METHODS

### Resource availability

#### Lead contact and materials availability

Further information and requests for resources and reagents should be directed to and will be fulfilled by the lead contact, Philippe Pierre mail to.

#### Data availability

scRNA seq data are available under submission number GSE277164 at https://www.ncbi.nlm.nih.gov/geo/query/acc.cgi?acc=GSE277164. Any additional information required to analyze the data reported in this paper is available from the lead contact upon request.

## EXPERIMENTAL MODEL AND SUBJECT DETAILS

### Cell lines

CAL-1 cells (gift from Dr. Takahiro Maeda, Nagasaki University, Japan) were grown in RPMI 1640 medium with 2 mM L-glutamine (Gibco, Thermo Fisher Scientific, Waltham, MA, USA), supplemented with 10% heat-inactivated foetal bovine serum (FBS, Sigma-Aldrich, St. Louis, MO, USA), 1x non-essential amino acids, 10 mM HEPES, and 1 mM sodium pyruvate (all from Gibco). For each experiment, cells bellow passage 15 were plated 16 h before at 1x10^6^ cells/mL in complete medium with 1% FBS. Cells were cultured in a humified atmosphere at 37°C with 5% CO_2_ and were regularly tested for mycoplasma contamination. Cells were stimulated with 1μg/ml of purified SubAB or mutant toxin and 1μM^18^ or the TLR7 agonist CL307^28^.

### Primary cell purification

PBMCs were isolated from healthy blood donors’ buffy coats (Établissement Français du Sang (EFS), Marseille, FR) by density gradient centrifugation, using Ficoll-Paque Plus (Cytiva, Marlborough, MA, USA). Briefly, the buffy coat blood was diluted 1:1 in RPMI 1640 medium and overlaid onto Ficoll-Paque Plus density gradient media, and cells separated by centrifugation (2,200 rpm for 30 min, no breaks). The mononuclear cell layer was transferred into a new tube and washed with RPMI 1640 medium at 1,900 rpm for 5 minutes. The final PBMCs pellet was resuspended at the desired concentration for downstream use.

pDCs were purified from the PBMCs of healthy donors (isolated as described above) using a negative selection MACS beads-based assay (Plasmacytoid Dendritic Cell Isolation Kit II human, Miltenyi Biotec, Bergisch Gladbach, GER), following manufacturer’s instructions. pDCs isolated by this procedure were > 80% pure and their viability was > 95%. Healthy pDCs were cultured in RPMI 1640 medium with 2 mM L-glutamine supplemented with 10% heat-inactivated human serum (Sigma-Aldrich), 1 mM penicillin/streptomycin (Gibco) and in the presence of IL-3 (1 ng/mL, PeproTech, London, UK).

## METHOD DETAILS

### Single-cell RNA sequencing (scRNA-seq) analyses

Single cell transcriptomics was performed to define, at the molecular level, the impact of the SubAB toxin on human immune cells. scRNA-seq experiments were performed at the “Labtech SingleCell@Imagine” on PBMCs isolated from healthy blood donors’ buffy coats. The scRNA-seq libraries were generated using Chromium Single Cell 30 Library & Gel Bead Kit v.3 (10x Genomics) according to the manufacturer’s protocol. Briefly, cells were counted, diluted at 1,000 cells/mL in PBS and 20,000 cells were loaded in the 10x Chromium Controller to generate single-cell gel-beads in emulsion. After reverse transcription, gel-beads in emulsion were disrupted. Barcoded complementary-DNA was isolated and amplified by PCR. Following fragmentation, end repair, and A-tailing, sample indexes were added during index PCR. The purified libraries were sequenced on a Novaseq 6000 (Illumina) with 28 cycles of read 1, 8 cycles of i7 index and 91 cycles of read 2.

### Bioinformatics analysis of scRNA-seq data

Sequencing reads were demultiplexed and aligned to the human reference genome (GRCh38-2020-A), using the CellRanger Pipeline v7.1.0. Unfiltered RNA UMI counts were loaded into Seurat v5.01 for quality control^86^, data integration and downstream analyses. Apoptotic cells and empty sequencing capsules were excluded by filtering out cells with less than 500 features or a mitochondrial content higher than 20%. Data from each sample were SCT normalised, before batch correction using Seurat’s FindIntegratedAnchors. For computational efficiency, anchors for integration were determined using all control samples as reference and patient samples were projected onto the integrated controls space. On this integrated dataset, the principal component analysis was computed on the 3000 most variable genes. UMAP was carried out using the 20 most significant principal components (PCs), and community detection was performed using the graph-based modularity-optimization Louvain algorithm from Seurat’s FindClusters function with a 1.4 resolution. Cell types of labels were assigned to resulting clusters based on a manually curated list of marker genes as well as previously defined signatures of the well-known PBMCs subtypes. The gene signature scores were calculated using Ucell v2.6^88^. After extraction and re-clustering of high-quality cells, differential expression was performed separately on all PBMCs, monocytes/DCs, T cells or B cells. Differential expression testing was conducted using the FindMarkers function of Seurat on the SCT assay with default parameters. Genes with adjusted p values %0.05 were selected as significant. Pathways analysis was performed with EnrichR^30^ on data found under GSE277164. Heat maps were generated using the Morpheus software (https://software.broadinstitute.org/morpheus) based in P-values of top significantly enriched pathways found in MSigDB 2020 by EnrichR^30^.

### CRISPR/Cas9 knockout

Gene inactivation was performed with CRISPR/Cas9 technology in an inducible Cas9 expressing CAL-1 cell line, which constitutively expressed green fluorescent protein (GFP) (CAL-1-Cas9-GFP). The genomic target sequences were searched by accessing the GeneArt CRISPR Search and Design (thermofisher.com/crisprdesign) or based in the literature. Guide RNAs (gRNA) were generated using the GeneArt Precision gRNA synthesis kit (Invitrogen). All gRNA target sequences are listed in **Table S3**. To induce Cas9 expression, 6.25x10^5^ CAL-1-Cas9-GFP cells were seeded in a 24-well plate in the presence of 2 μg/ml doxycycline (Sigma-Aldrich) for 48 h before gRNA transfection. For gRNA delivery, 5 μg of gRNA for the gene of interest and gRNA for GFP gene were co-transfected using electroporation in 100 μL of non-supplemented RPMI 1640 medium at room temperature (RT) with the following settings: one pulse of 350 V (875 V/cm), 950 μF and 25 Ω in a 4 mm cuvette (MBP Molecular BioProducts, Toronto, CA), by BTX ECM electroporation system (BTX, Biochrom, Cambridge, UK). The efficiency of transfection was assessed based on the decrease of GFP expression by flow cytometry using a BD Accuri C6 cytometer (BD Biosciences, San Jose, CA, USA). GFP-negative clones were selected by limiting dilutions and individual clones were analysed by Western-Blotting and conventional PCR. Genomic DNA was isolated using NZY Tissue gDNA Isolation Kit (NZYTech, Lisbon, PT) and the target site was amplified by PCR using the AmpliTaq Gold™ 360 Master Mix (Thermo Fisher Scientific). The primer sequences are presented in **Table S4**. Sanger sequencing was used to characterize indels. The sequencing results were analyzed using Geneious Pro 4.8.2 (Biomatters, Auckland, NZ).

### Quantitative PCR

Total mRNA from cells was purified using the RNeasy Mini Kit (Qiagen, Venlo, NL). For cDNA synthesis, 500 ng of total RNA were subjected to reverse transcription using SuperScript II (Invitrogen, Thermo Fisher Scientific). Quantitative PCR was performed using 2x SYBR Green qPCR Master Mix (Low Rox; Selleckchem, Houston, TX, USA) with 7500 Real-Time PCR system (Applied Biosystems, Thermo Fisher Scientific). The relative amount of each transcript was determined after normalization to the internal housekeeping gene expression (*gapdh*). A list of primers can be found in **Table S4.**

### Translation intensity measurement

Protein synthesis was monitored using puromycin labeling and performed as previously described ^46^. Cells were incubated with puromycin (Sigma-Aldrich) at 1 µg/ml for 15 min at 37°C and 5% CO_2_. After puromycin incorporation the cells were harvested, washed in PBS and incubated for 30 min at 4°C in the dark with Live/Dead™ fixable green dead cell stain (Invitrogen) to allow dead cell exclusion. Cells were then fixed in 4% paraformaldehyde (PFA; Alfa Aesar, Haverhill, MA, USA) for 15 min at RT. For cell permeabilization, cells were incubated for 15 min in FACS buffer (PBS, 0.5% BSA (NZYTech), 0.01% sodium azide (Sigma-Aldrich), 0.1% saponin (Alfa-Aesar)) at 4°C. Cells were then stained with a mouse monoclonal anti-puromycin-AF647 (Clone 12D10; Merck Millipore, Burlington, MA, USA). After 1 h of incubation, cells were washed, resuspended in PBS and dissociated/filtered using a 40 μm cell strainer. Samples were acquired in a BD Accuri C6 Flow Cytometer (BD Biosciences, San Jose, CA, USA). BD Accuri S6 software (BD Biosciences) was used for analyzes.

### Flow cytometry

Cell suspensions were washed and incubated with a cocktail of coupled specific antibodies for cell surface markers in FACS buffer (PBS, 1% FBS and 2 mM EDTA) for 30 min at 4°C. For intracellular staining, cells were next fixed with BD phosflow fix buffer I during 10 min at RT and washed with 10% Perm/wash Buffer I 1x (BD Biosciences). Permeabilized cells were blocked for 10 min with 10% Perm/Wash buffer 1x, 10% FCS, before staining with primary antibodies. When the primary antibody was not coupled, cells were washed after and blocked during 10 min with Perm/Wash buffer 1x, 10% FCS, and 10% of serum from the species where the secondary antibody was produced. Then, the incubation with the secondary antibody was performed at 4°C for 30 min. The events were collected in a FACS LSRUVII (BD Biosciences) and the events acquired by FACS DIVA software (BD Biosciences) and analyzed by FlowJow or OMICs software. The list of antibodies used can be found on Key resources table.

For the analysis of SubAB-treated PBMC samples using spectral flow cytometry, 3 healthy donor samples were used with treatments done in duplicates. For surface staining, cells were pre-incubated with 20µL of a mix of anti-γδTCR antibody (1:50 dilution; Cytek) and FcR blocking reagent (1:50 dilution; Miltenyi Biotech) diluted in PBS 1x for 10 min prior to a 20 min staining with 30µL of a second mix with the surface antibodies listed on Key resources table. Afterwards, cells were fixed/permeabilized for 30 min with Foxp3 fixation/permeabilization reagent 1x (Invitrogen), washed with Foxp3 Permeabilization buffer 1X (Invitrogen) and stained intracellularly with anti-Foxp3 (1:50 dilution; BD Biosciences) and anti-perforin (1:50 dilution; Biolegend) antibodies diluted in Foxp3 Permeabilization buffer 1x for 1 h at 4°C. Cells were then washed and resuspended in FACS buffer before acquiring in a 5L Aurora Cytek spectral analyzer (BD Biosciences). FACS data was analyzed using the OMIQ platform (Dotmatics, Boston, MA, USA). Data quality control and cleanup was done with flowAI and 3x10^5^ cells from the Singlets/Live/CD45+ gate from each sample/condition were concatenated and further analyzed using unsupervised clustering analysis with UMAP for dimensionality reduction and FlowSOM as a clustering algorithm to profile the different cell subpopulations.

### Immunoblotting

Cell pellets were lysed with radioimmunoprecipitation assay (RIPA) buffer (25 mM Tris-HCl pH 7.6, 150 mM NaCl, 1% NP-40; 1% sodium deoxycholate, and 0.1% SDS) supplemented with Pierce Protease Inhibitor Tablets (Thermo Fisher Scientific), phosphatase inhibitors – sodium fluoride (NaF, 50 mM) and sodium orthovanadate (Na_3_VO_4_, 0.2 mM), both from Sigma-Aldrich – and a proteosome inhibitor (MG132, 5 µM; CNSpharma, Houston, TX, USA). Protein quantification was performed using the bicinchoninic acid (BCA) protein assay (Pierce, Thermo Fisher Scientific). A total of 20-30 µg of soluble proteins were run in SDS-polyacrylamide gels. Gel transfer was performed using PVDF membranes (Millipore) and the membrane were incubated in blocking solution [Tris-buffered saline with 0.05% Tween 20 (TBS-T, Sigma Aldrich) and 5% BSA (NZYTech)] prior antibody binding and chemiluminescence detection. Image Lab Software (Bio-Rad, Hercules, CA, USA) was used for quantification of the protein bands. Antibodies used for immunoblotting are listed on Key resources table.

### Immunofluorescence

Cells were harvested and seeded on 12-mm Alcian blue (Sigma-Aldrich)-pretreated coverslips. The coverslips were then kept at 37°C for 10 min and fixed with 3.7% PFA for 10 min at RT. Cells were permeabilized (0.1 % Triton X-100, 5% FBS, 100 mM glycine, 1% PBS) for 15 min at RT. Incubation with primary antibody anti-G3BP1 (Santa Cruz Biotechnology, Dallas, TX, USA) was performed overnight at 4°C, followed by the anti-rabbit IgG-Alexa Fluor 647 secondary antibody (Invitrogen). Coverslips were mounted in ProLong Gold DAPI (Invitrogen). Images were visualized in a Zeiss LSM780 confocal microscope (Zeiss, Oberkochen, GER) using 63x objective and accompanying imaging software^89^. Details of the antibodies used are listed on Key resources table.

### Cytokines quantification by ELISA

Human IFN-β was quantified in cell supernatants using a commercially available ELISA kit from InvivoGen - LumiKin Xpress hIFN-β 2.0. The assays were performed according to the manufacturer’s instructions.

### SDD-AGE

Subcellular fractionation was performed using a Mitochondrial Isolation Kit for cultured cells (Thermo Fisher Scientific), following the manufacturer’s instructions. The P5 fractions containing the crude mitochondria were used to evaluate the ability of SubAB to induce the formation of MAVS aggregates by SDD-AGE.

Crude mitochondria (P5 fraction) were isolated from CAL-1 cells, resuspended in the sample buffer (0.5x TBE, 10% glycerol, 2% SDS and 0.0025% bromophenol blue) and loaded onto a vertical 1.5% agarose gel. The gels were run in migration buffer (1x TBE and 0.1% SDS) for 30 min with a constant voltage of 100 V at 4°C. Immunoblotting was then performed as described above.

## QUANTIFICATION AND STATISTICAL ANALYSIS

### Statistical analysis

Statistical analysis was performed using GraphPad Prism Software Version 9.0.0 (GraphPad Software, Inc., La Jolla, CA, USA). The Shapiro-Wilk test was used to assess data normality. The most appropriate statistical test was then chosen according to each set of data as indicated in figure legends. Data is presented as mean ± standard deviation (SD). \**p* ≤ 0.05; \*\**p* ≤ 0.01; \*\*\**p* ≤ 0.001; \*\*\*\**p* ≤ 0.0001.

## SUPPLEMENTAL INFORMATION

Document S1. Figures S1-S8, Tables S2-S4 and supplemental references. Table S1. Excel file containing additional pathway analysis related to Figure 2.

### Lists of the biological pathways most impacted by SubAB and CL307 treatment in different PBMC subsets

The most significant GO pathways (top 2 to 4 ranked by P-values) identified by scRNA sequencing are shown presented by cell type and stimuli. These gene signature pathways are representing transcription factors or signaling networks (TRUST TF2019 and MSigDB 2020) and identified genes present in the signatures are listed. **(A)** Monocytes, **(B)** B cells, **(C)** CD4^+^ T cells, **(D)** CD8^+^ T cells. **(E)** γδ T cells, **(F)** NK cells, **(G)** cDC, **(H)** Known STAT1-dependent genes (extracted from GSE48970 data set) and present in the IFN-a MSigDB 2020 signature.

## ACKNOWLEDGEMENTS

Work developed within the scope of iBiMED – Institute of Biomedicine (UIDB/04501/2020 and UIDP/04501/2020) and CICECO – Aveiro Institute of Materials UIDB/50011/2020 (DOI 10.54499/UIDB/50011/2020), UIDP/50011/2020 (DOI 10.54499/UIDP/50011/2020) & LA/P/0006/2020 (DOI 10.54499/LA/P/0006/2020) and the project with the reference 2022.03217.PTDC (DOI 10.54499/2022.03217.PTDC), financially supported by national funds (OE), through Fundação para a Ciência e a Tecnologia (FCT)/MCTES. The P.P laboratory is Equipe FRM sponsored by the grant DEQ20180339212, and grants from the Institut National du Cancer (INCA) PLBIO-2021 “Translec”, Fondation ARC “PGA 2021-2025-CHARP” and Agence Nationale de la Recherche (ANR) AAPG2021-STIM. We thank the “Shanghai 1000 talents” program and the Maratona da Saúde for their support. B.H.F., P.A. and F.L-P were supported by FCT through an individual grants SFRH/BD/144706/2019, SFRH/BD/138336/2018 and COVID/BD/153248/2023, and DOI 10.54499/2020.08489. Y.L and L.Z were supported by grant C24185. This work has been supported by CENTURI postdoc funding P.G.G and Horizon2020 Transcan 2021-227-TALETE French National Cancer Institute (INCa) to RJA. Image acquisition was performed in the LiM facility of iBiMED, a node of PPBI (Portuguese Platform of BioImaging): POCI-01-0145-FEDER-022122. We thank the CIML cytometry and cell sorting core facility for its support.

## Author contributions

D.B., B.H.F., F.L-P., P.A., A.M., M.D.M., F.C, P.G-G, J.G., M.R., M.C., M.L., L.G., L.Z performed research. A.W.P, J.C.P, M.N. B.S., S.R., Y.L. provided key reagents. D.B, B.F., B.N., R.J.A., F.R-L, M.M., E.G., C.A. and P.P. designed research and analysed data. D.B., B.F., C.A. and P.P. wrote the paper.

## Declaration of generative AI and AI-assisted technologies

During the preparation of this work, the author(s) used CHAT-GPT to simplify, correct English and avoid repetition in the text of this manuscript. After using this tool, the authors reviewed and edited the content as needed and take full responsibility for the content of the publication

## References

1. Lin, J.H., Walter, P., and Yen, T.S. (2008). Endoplasmic reticulum stress in disease pathogenesis. Annu Rev Pathol 3, 399–425. 10.1146/annurev.pathmechdis.3.121806.151434.

2. Walter, P., and Ron, D. (2011). The unfolded protein response: from stress pathway to homeostatic regulation. Science 334, 1081–1086. 10.1126/science.1209038.

3. Hetz, C., Zhang, K., and Kaufman, R.J. (2020). Mechanisms, regulation and functions of the unfolded protein response. Nat Rev Mol Cell Biol 21, 421–438. 10.1038/s41580-020-0250-z.

4. Ron, D., and Walter, P. (2007). Signal integration in the endoplasmic reticulum unfolded protein response. Nat Rev Mol Cell Biol 8, 519–529. 10.1038/nrm2199.

5. Han, J., Back, S.H., Hur, J., Lin, Y.H., Gildersleeve, R., Shan, J., Yuan, C.L., Krokowski, D., Wang, S., Hatzoglou, M., et al. (2013). ER-stress-induced transcriptional regulation increases protein synthesis leading to cell death. Nat Cell Biol 15, 481–490. 10.1038/ncb2738.

6. Costa-Mattioli, M., and Walter, P. (2020). The integrated stress response: From mechanism to disease. Science 368. 10.1126/science.aat5314.

7. Reimold, A.M., Iwakoshi, N.N., Manis, J., Vallabhajosyula, P., Szomolanyi-Tsuda, E., Gravallese, E.M., Friend, D., Grusby, M.J., Alt, F., and Glimcher, L.H. (2001). Plasma cell differentiation requires the transcription factor XBP-1. Nature 412, 300–307. 10.1038/35085509.

8. Iwakoshi, N.N., Pypaert, M., and Glimcher, L.H. (2007). The transcription factor XBP-1 is essential for the development and survival of dendritic cells. J Exp Med 204, 2267–2275. 10.1084/jem.20070525.

9. Todd, D.J., McHeyzer-Williams, L.J., Kowal, C., Lee, A.H., Volpe, B.T., Diamond, B., McHeyzer-Williams, M.G., and Glimcher, L.H. (2009). XBP1 governs late events in plasma cell differentiation and is not required for antigen-specific memory B cell development. J Exp Med 206, 2151–2159. 10.1084/jem.20090738.

10. Bettigole, S.E., and Glimcher, L.H. (2015). Endoplasmic reticulum stress in immunity. Annu Rev Immunol 33, 107–138. 10.1146/annurev-immunol-032414-112116.

11. Reizis, B., Colonna, M., Trinchieri, G., Barrat, F., and Gilliet, M. (2011). Plasmacytoid dendritic cells: one-trick ponies or workhorses of the immune system? Nat Rev Immunol 11, 558–565. 10.1038/nri3027.

12. Sprooten, J., and Garg, A.D. (2020). Type I interferons and endoplasmic reticulum stress in health and disease. Int Rev Cell Mol Biol 350, 63–118. 10.1016/bs.ircmb.2019.10.004.

13. Ishikawa, H., Ma, Z., and Barber, G.N. (2009). STING regulates intracellular DNA-mediated, type I interferon-dependent innate immunity. Nature 461, 788–792. 10.1038/nature08476.

14. Ritchie, C., Carozza, J.A., and Li, L. (2022). Biochemistry, Cell Biology, and Pathophysiology of the Innate Immune cGAS-cGAMP-STING Pathway. Annu Rev Biochem 91, 599–628. 10.1146/annurev-biochem-040320-101629.

15. Moretti, J., Roy, S., Bozec, D., Martinez, J., Chapman, J.R., Ueberheide, B., Lamming, D.W., Chen, Z.J., Horng, T., Yeretssian, G., et al. (2017). STING Senses Microbial Viability to Orchestrate Stress-Mediated Autophagy of the Endoplasmic Reticulum. Cell 171, 809–823 e813. 10.1016/j.cell.2017.09.034.

16. Wu, J., Chen, Y.J., Dobbs, N., Sakai, T., Liou, J., Miner, J.J., and Yan, N. (2019). STING-mediated disruption of calcium homeostasis chronically activates ER stress and primes T cell death. J Exp Med 216, 867–883. 10.1084/jem.20182192.

17. Zhang, D., Liu, Y., Zhu, Y., Zhang, Q., Guan, H., Liu, S., Chen, S., Mei, C., Chen, C., Liao, Z., et al. (2022). A non-canonical cGAS-STING-PERK pathway facilitates the translational program critical for senescence and organ fibrosis. Nat Cell Biol 24, 766–782. 10.1038/s41556-022-00894-z.

18. Paton, A.W., Srimanote, P., Talbot, U.M., Wang, H., and Paton, J.C. (2004). A new family of potent AB(5) cytotoxins produced by Shiga toxigenic Escherichia coli. J Exp Med 200, 35–46. 10.1084/jem.20040392.

19. Smith, R.D., Willett, R., Kudlyk, T., Pokrovskaya, I., Paton, A.W., Paton, J.C., and Lupashin, V.V. (2009). The COG complex, Rab6 and COPI define a novel Golgi retrograde trafficking pathway that is exploited by SubAB toxin. Traffic 10, 1502–1517. 10.1111/j.1600-0854.2009.00965.x.

20. Chong, D.C., Paton, J.C., Thorpe, C.M., and Paton, A.W. (2008). Clathrin-dependent trafficking of subtilase cytotoxin, a novel AB5 toxin that targets the endoplasmic reticulum chaperone BiP. Cell Microbiol 10, 795–806. 10.1111/j.1462-5822.2007.01085.x.

21. Byres, E., Paton, A.W., Paton, J.C., Lofling, J.C., Smith, D.F., Wilce, M.C., Talbot, U.M., Chong, D.C., Yu, H., Huang, S., et al. (2008). Incorporation of a non-human glycan mediates human susceptibility to a bacterial toxin. Nature 456, 648–652. 10.1038/nature07428.

22. Paton, A.W., Beddoe, T., Thorpe, C.M., Whisstock, J.C., Wilce, M.C., Rossjohn, J., Talbot, U.M., and Paton, J.C. (2006). AB5 subtilase cytotoxin inactivates the endoplasmic reticulum chaperone BiP. Nature 443, 548–552. 10.1038/nature05124.

23. Hu, C.C., Dougan, S.K., Winter, S.V., Paton, A.W., Paton, J.C., and Ploegh, H.L. (2009). Subtilase cytotoxin cleaves newly synthesized BiP and blocks antibody secretion in B lymphocytes. J Exp Med 206, 2429–2440. 10.1084/jem.20090782.

24. Harama, D., Koyama, K., Mukai, M., Shimokawa, N., Miyata, M., Nakamura, Y., Ohnuma, Y., Ogawa, H., Matsuoka, S., Paton, A.W., et al. (2009). A subcytotoxic dose of subtilase cytotoxin prevents lipopolysaccharide-induced inflammatory responses, depending on its capacity to induce the unfolded protein response. J Immunol 183, 1368–1374. 10.4049/jimmunol.0804066.

25. Nakajima, S., Saito, Y., Takahashi, S., Hiramatsu, N., Kato, H., Johno, H., Yao, J., Paton, A.W., Paton, J.C., and Kitamura, M. (2010). Anti-inflammatory subtilase cytotoxin up-regulates A20 through the unfolded protein response. Biochem Biophys Res Commun 397, 176–180. 10.1016/j.bbrc.2010.05.069.

26. Martinon, F., Chen, X., Lee, A.H., and Glimcher, L.H. (2010). TLR activation of the transcription factor XBP1 regulates innate immune responses in macrophages. Nat Immunol 11, 411–418. 10.1038/ni.1857.

27. Qian, T., Huo, B., Deng, X., Song, X., Jiang, Y., Yang, J., and Hao, F. (2023). Decreased TAX1BP1 participates in systemic lupus erythematosus by regulating monocyte/macrophage function. Int Immunol 35, 483–495. 10.1093/intimm/dxad027.

28. Gutjahr, A., Papagno, L., Nicoli, F., Lamoureux, A., Vernejoul, F., Lioux, T., Gostick, E., Price, D.A., Tiraby, G., Perouzel, E., et al. (2017). Cutting Edge: A Dual TLR2 and TLR7 Ligand Induces Highly Potent Humoral and Cell-Mediated Immune Responses. J Immunol 198, 4205–4209. 10.4049/jimmunol.1602131.

29. Kapellos, T.S., Bonaguro, L., Gemund, I., Reusch, N., Saglam, A., Hinkley, E.R., and Schultze, J.L. (2019). Human Monocyte Subsets and Phenotypes in Major Chronic Inflammatory Diseases. Front Immunol 10, 2035. 10.3389/fimmu.2019.02035.

30. de Cevins, C., Delage, L., Batignes, M., Riller, Q., Luka, M., Remaury, A., Sorin, B., Fali, T., Masson, C., Hoareau, B., et al. (2023). Single-cell RNA-sequencing of PBMCs from SAVI patients reveals disease-associated monocytes with elevated integrated stress response. Cell Rep Med 4, 101333. 10.1016/j.xcrm.2023.101333.

31. Kuleshov, M.V., Jones, M.R., Rouillard, A.D., Fernandez, N.F., Duan, Q., Wang, Z., Koplev, S., Jenkins, S.L., Jagodnik, K.M., Lachmann, A., et al. (2016). Enrichr: a comprehensive gene set enrichment analysis web server 2016 update. Nucleic Acids Res 44, W90–97. 10.1093/nar/gkw377.

32. Ancuta, P., Liu, K.Y., Misra, V., Wacleche, V.S., Gosselin, A., Zhou, X., and Gabuzda, D. (2009). Transcriptional profiling reveals developmental relationship and distinct biological functions of CD16+ and CD16-monocyte subsets. BMC Genomics 10, 403. 10.1186/1471-2164-10-403.

33. Subramanian, A., Tamayo, P., Mootha, V.K., Mukherjee, S., Ebert, B.L., Gillette, M.A., Paulovich, A., Pomeroy, S.L., Golub, T.R., Lander, E.S., and Mesirov, J.P. (2005). Gene set enrichment analysis: a knowledge-based approach for interpreting genome-wide expression profiles. Proc Natl Acad Sci U S A 102, 15545–15550. 10.1073/pnas.0506580102.

34. Goodall, J.C., Wu, C., Zhang, Y., McNeill, L., Ellis, L., Saudek, V., and Gaston, J.S. (2010). Endoplasmic reticulum stress-induced transcription factor, CHOP, is crucial for dendritic cell IL-23 expression. Proc Natl Acad Sci U S A 107, 17698–17703. 10.1073/pnas.1011736107.

35. Liu, Y.J. (2005). IPC: professional type 1 interferon-producing cells and plasmacytoid dendritic cell precursors. Annu Rev Immunol 23, 275–306. 10.1146/annurev.immunol.23.021704.115633.

36. Clavarino, G., Claudio, N., Dalet, A., Terawaki, S., Couderc, T., Chasson, L., Ceppi, M., Schmidt, E.K., Wenger, T., Lecuit, M., et al. (2012). Protein phosphatase 1 subunit Ppp1r15a/GADD34 regulates cytokine production in polyinosinic:polycytidylic acid-stimulated dendritic cells. Proc Natl Acad Sci U S A 109, 3006–3011. 10.1073/pnas.1104491109.

37. Sidrauski, C., McGeachy, A.M., Ingolia, N.T., and Walter, P. (2015). The small molecule ISRIB reverses the effects of eIF2alpha phosphorylation on translation and stress granule assembly. Elife 4. 10.7554/eLife.05033.

38. Combes, A., Camosseto, V., N’Guessan, P., Arguello, R.J., Mussard, J., Caux, C., Bendriss-Vermare, N., Pierre, P., and Gatti, E. (2017). BAD-LAMP controls TLR9 trafficking and signalling in human plasmacytoid dendritic cells. Nat Commun 8, 913. 10.1038/s41467-017-00695-1.

39. Narita, M., Watanabe, N., Yamahira, A., Hashimoto, S., Tochiki, N., Saitoh, A., Kaji, M., Nakamura, T., Furukawa, T., Toba, K., et al. (2009). A leukemic plasmacytoid dendritic cell line, PMDC05, with the ability to secrete IFN-alpha by stimulation via Toll-like receptors and present antigens to naive T cells. Leuk Res 33, 1224–1232. 10.1016/j.leukres.2009.03.047.

40. Steinhagen, F., Meyer, C., Tross, D., Gursel, M., Maeda, T., Klaschik, S., and Klinman, D.M. (2012). Activation of type I interferon-dependent genes characterizes the “core response” induced by CpG DNA. J Leukoc Biol 92, 775–785. 10.1189/jlb.1011522.

41. Carmona-Saez, P., Varela, N., Luque, M.J., Toro-Dominguez, D., Martorell-Marugan, J., Alarcon-Riquelme, M.E., and Maranon, C. (2017). Metagene projection characterizes GEN2.2 and CAL-1 as relevant human plasmacytoid dendritic cell models. Bioinformatics 33, 3691–3695. 10.1093/bioinformatics/btx502.

42. Axten, J.M., Romeril, S.P., Shu, A., Ralph, J., Medina, J.R., Feng, Y., Li, W.H., Grant, S.W., Heerding, D.A., Minthorn, E., et al. (2013). Discovery of GSK2656157: An Optimized PERK Inhibitor Selected for Preclinical Development. ACS Med Chem Lett 4, 964–968. 10.1021/ml400228e.

43. Mendes, A., Gigan, J.P., Rodriguez Rodrigues, C., Choteau, S.A., Sanseau, D., Barros, D., Almeida, C., Camosseto, V., Chasson, L., Paton, A.W., et al. (2021). Proteostasis in dendritic cells is controlled by the PERK signaling axis independently of ATF4. Life Sci Alliance 4. 10.26508/lsa.202000865.

44. Garcia-Gonzalez, P., Fernandez, D., Gutierrez, D., Parra-Cordero, M., and Osorio, F. (2022). Human cDC1s display constitutive activation of the UPR sensor IRE1. Eur J Immunol 52, 1069–1076. 10.1002/eji.202149774.

45. Yahiro, K., Ogura, K., Tsutsuki, H., Iyoda, S., Ohnishi, M., and Moss, J. (2021). A novel endoplasmic stress mediator, Kelch domain containing 7B (KLHDC7B), increased Harakiri (HRK) in the SubAB-induced apoptosis signaling pathway. Cell Death Discov 7, 360. 10.1038/s41420-021-00753-0.

46. Schmidt, E.K., Clavarino, G., Ceppi, M., and Pierre, P. (2009). SUnSET, a nonradioactive method to monitor protein synthesis. Nat Methods 6, 275–277. 10.1038/nmeth.1314.

47. Shoulders, M.D., Ryno, L.M., Genereux, J.C., Moresco, J.J., Tu, P.G., Wu, C., Yates, J.R., 3rd, Su, A.I., Kelly, J.W., and Wiseman, R.L. (2013). Stress-independent activation of XBP1s and/or ATF6 reveals three functionally diverse ER proteostasis environments. Cell Rep 3, 1279–1292. 10.1016/j.celrep.2013.03.024.

48. Harding, H.P., Zhang, Y., Scheuner, D., Chen, J.J., Kaufman, R.J., and Ron, D. (2009). Ppp1r15 gene knockout reveals an essential role for translation initiation factor 2 alpha (eIF2alpha) dephosphorylation in mammalian development. Proc Natl Acad Sci U S A 106, 1832–1837. 10.1073/pnas.0809632106.

49. Ivanov, P., Kedersha, N., and Anderson, P. (2018). Stress Granules and Processing Bodies in Translational Control. Cold Spring Harb Perspect Biol. 10.1101/cshperspect.a032813.

50. Winkler, R., Gillis, E., Lasman, L., Safra, M., Geula, S., Soyris, C., Nachshon, A., Tai-Schmiedel, J., Friedman, N., Le-Trilling, V.T.K., et al. (2019). m(6)A modification controls the innate immune response to infection by targeting type I interferons. Nat Immunol 20, 173–182. 10.1038/s41590-018-0275-z.

51. Cerezo, M., Lehraiki, A., Millet, A., Rouaud, F., Plaisant, M., Jaune, E., Botton, T., Ronco, C., Abbe, P., Amdouni, H., et al. (2016). Compounds Triggering ER Stress Exert Anti-Melanoma Effects and Overcome BRAF Inhibitor Resistance. Cancer Cell 29, 805–819. 10.1016/j.ccell.2016.04.013.

52. Tan, X., Sun, L., Chen, J., and Chen, Z.J. (2018). Detection of Microbial Infections Through Innate Immune Sensing of Nucleic Acids. Annu Rev Microbiol 72, 447–478. 10.1146/annurev-micro-102215-095605.

53. Ablasser, A., and Hur, S. (2020). Regulation of cGAS- and RLR-mediated immunity to nucleic acids. Nat Immunol 21, 17–29. 10.1038/s41590-019-0556-1.

54. Liu, S., Cai, X., Wu, J., Cong, Q., Chen, X., Li, T., Du, F., Ren, J., Wu, Y.T., Grishin, N.V., and Chen, Z.J. (2015). Phosphorylation of innate immune adaptor proteins MAVS, STING, and TRIF induces IRF3 activation. Science 347, aaa2630. 10.1126/science.aaa2630.

55. Balka, K.R., Louis, C., Saunders, T.L., Smith, A.M., Calleja, D.J., D’Silva, D.B., Moghaddas, F., Tailler, M., Lawlor, K.E., Zhan, Y., et al. (2020). TBK1 and IKKepsilon Act Redundantly to Mediate STING-Induced NF-kappaB Responses in Myeloid Cells. Cell Rep 31, 107492. 10.1016/j.celrep.2020.03.056.

56. Clark, K., Peggie, M., Plater, L., Sorcek, R.J., Young, E.R., Madwed, J.B., Hough, J., McIver, E.G., and Cohen, P. (2011). Novel cross-talk within the IKK family controls innate immunity. Biochem J 434, 93–104. 10.1042/BJ20101701.

57. Hou, F., Sun, L., Zheng, H., Skaug, B., Jiang, Q.X., and Chen, Z.J. (2011). MAVS forms functional prion-like aggregates to activate and propagate antiviral innate immune response. Cell 146, 448–461. 10.1016/j.cell.2011.06.041.

58. Abdel-Nour, M., Carneiro, L.A.M., Downey, J., Tsalikis, J., Outlioua, A., Prescott, D., Da Costa, L.S., Hovingh, E.S., Farahvash, A., Gaudet, R.G., et al. (2019). The heme-regulated inhibitor is a cytosolic sensor of protein misfolding that controls innate immune signaling. Science 365. 10.1126/science.aaw4144.

59. Qi, N., Shi, Y., Zhang, R., Zhu, W., Yuan, B., Li, X., Wang, C., Zhang, X., and Hou, F. (2017). Multiple truncated isoforms of MAVS prevent its spontaneous aggregation in antiviral innate immune signalling. Nat Commun 8, 15676. 10.1038/ncomms15676.

60. Shi, Y., Wu, J., Zhong, T., Zhu, W., She, G., Tang, H., Du, W., Ye, B.C., and Qi, N. (2020). Upstream ORFs Prevent MAVS Spontaneous Aggregation and Regulate Innate Immune Homeostasis. iScience 23, 101059. 10.1016/j.isci.2020.101059.

61. Barber, G.N. (2015). STING: infection, inflammation and cancer. Nat Rev Immunol 15, 760–770. 10.1038/nri3921.

62. Jeremiah, N., Neven, B., Gentili, M., Callebaut, I., Maschalidi, S., Stolzenberg, M.C., Goudin, N., Fremond, M.L., Nitschke, P., Molina, T.J., et al. (2014). Inherited STING-activating mutation underlies a familial inflammatory syndrome with lupus-like manifestations. J Clin Invest 124, 5516–5520. 10.1172/JCI79100.

63. Steiner, A., Hrovat-Schaale, K., Prigione, I., Yu, C.H., Laohamonthonkul, P., Harapas, C.R., Low, R.R.J., De Nardo, D., Dagley, L.F., Mlodzianoski, M.J., et al. (2022). Deficiency in coatomer complex I causes aberrant activation of STING signalling. Nat Commun 13, 2321. 10.1038/s41467-022-29946-6.

64. Konno, H., Konno, K., and Barber, G.N. (2013). Cyclic dinucleotides trigger ULK1 (ATG1) phosphorylation of STING to prevent sustained innate immune signaling. Cell 155, 688–698. 10.1016/j.cell.2013.09.049.

65. Klionsky, D.J., Abdel-Aziz, A.K., Abdelfatah, S., Abdellatif, M., Abdoli, A., Abel, S., Abeliovich, H., Abildgaard, M.H., Abudu, Y.P., Acevedo-Arozena, A., et al. (2021). Guidelines for the use and interpretation of assays for monitoring autophagy (4th edition)(1). Autophagy 17, 1–382. 10.1080/15548627.2020.1797280.

66. Johansen, T., and Lamark, T. (2011). Selective autophagy mediated by autophagic adapter proteins. Autophagy 7, 279–296.

67. Birgisdottir, A.B., Lamark, T., and Johansen, T. (2013). The LIR motif - crucial for selective autophagy. J Cell Sci 126, 3237–3247. 10.1242/jcs.126128.

68. Deretic, V. (2021). Autophagy in inflammation, infection, and immunometabolism. Immunity 54, 437–453. 10.1016/j.immuni.2021.01.018.

69. Sarraf, S.A., Shah, H.V., Kanfer, G., Pickrell, A.M., Holtzclaw, L.A., Ward, M.E., and Youle, R.J. (2020). Loss of TAX1BP1-Directed Autophagy Results in Protein Aggregate Accumulation in the Brain. Mol Cell 80, 779–795 e710. 10.1016/j.molcel.2020.10.041.

70. Choi, Y.B., Shembade, N., Parvatiyar, K., Balachandran, S., and Harhaj, E.W. (2017). TAX1BP1 Restrains Virus-Induced Apoptosis by Facilitating Itch-Mediated Degradation of the Mitochondrial Adaptor MAVS. Mol Cell Biol 37. 10.1128/MCB.00422-16.

71. Shembade, N., Pujari, R., Harhaj, N.S., Abbott, D.W., and Harhaj, E.W. (2011). The kinase IKKalpha inhibits activation of the transcription factor NF-kappaB by phosphorylating the regulatory molecule TAX1BP1. Nat Immunol 12, 834–843. 10.1038/ni.2066.

72. Verstrepen, L., Verhelst, K., Carpentier, I., and Beyaert, R. (2011). TAX1BP1, a ubiquitin-binding adaptor protein in innate immunity and beyond. Trends Biochem Sci 36, 347–354. 10.1016/j.tibs.2011.03.004.

73. Parvatiyar, K., Barber, G.N., and Harhaj, E.W. (2010). TAX1BP1 and A20 inhibit antiviral signaling by targeting TBK1-IKKi kinases. J Biol Chem 285, 14999–15009. 10.1074/jbc.M110.109819.

74. Yoshimura, A., Ito, M., Chikuma, S., Akanuma, T., and Nakatsukasa, H. (2018). Negative Regulation of Cytokine Signaling in Immunity. Cold Spring Harb Perspect Biol 10. 10.1101/cshperspect.a028571.

75. Bago, R., Malik, N., Munson, M.J., Prescott, A.R., Davies, P., Sommer, E., Shpiro, N., Ward, R., Cross, D., Ganley, I.G., and Alessi, D.R. (2014). Characterization of VPS34-IN1, a selective inhibitor of Vps34, reveals that the phosphatidylinositol 3-phosphate-binding SGK3 protein kinase is a downstream target of class III phosphoinositide 3-kinase. Biochem J 463, 413–427. 10.1042/BJ20140889.

76. Mejlvang, J., Olsvik, H., Svenning, S., Bruun, J.A., Abudu, Y.P., Larsen, K.B., Brech, A., Hansen, T.E., Brenne, H., Hansen, T., et al. (2018). Starvation induces rapid degradation of selective autophagy receptors by endosomal microautophagy. J Cell Biol 217, 3640–3655. 10.1083/jcb.201711002.

77. Wang, H., Paton, J.C., and Paton, A.W. (2007). Pathologic changes in mice induced by subtilase cytotoxin, a potent new Escherichia coli AB5 toxin that targets the endoplasmic reticulum. J Infect Dis 196, 1093–1101. 10.1086/521364.

78. Chimen, M., Yates, C.M., McGettrick, H.M., Ward, L.S., Harrison, M.J., Apta, B., Dib, L.H., Imhof, B.A., Harrison, P., Nash, G.B., and Rainger, G.E. (2017). Monocyte Subsets Coregulate Inflammatory Responses by Integrated Signaling through TNF and IL-6 at the Endothelial Cell Interface. J Immunol 198, 2834–2843. 10.4049/jimmunol.1601281.

79. Gentili, M., Liu, B., Papanastasiou, M., Dele-Oni, D., Schwartz, M.A., Carlson, R.J., Al’Khafaji, A.M., Krug, K., Brown, A., Doench, J.G., et al. (2023). ESCRT-dependent STING degradation inhibits steady-state and cGAMP-induced signalling. Nat Commun 14, 611. 10.1038/s41467-023-36132-9.

80. Woo, C.W., Cui, D., Arellano, J., Dorweiler, B., Harding, H., Fitzgerald, K.A., Ron, D., and Tabas, I. (2009). Adaptive suppression of the ATF4-CHOP branch of the unfolded protein response by toll-like receptor signalling. Nat Cell Biol 11, 1473–1480. 10.1038/ncb1996.

81. Arguello, R.J., Rodriguez Rodrigues, C., Gatti, E., and Pierre, P. (2015). Protein synthesis regulation, a pillar of strength for innate immunity? Curr Opin Immunol 32, 28–35. 10.1016/j.coi.2014.12.001.

82. Jiang, H.Y., Wek, S.A., McGrath, B.C., Lu, D., Hai, T., Harding, H.P., Wang, X., Ron, D., Cavener, D.R., and Wek, R.C. (2004). Activating transcription factor 3 is integral to the eukaryotic initiation factor 2 kinase stress response. Mol Cell Biol 24, 1365–1377.

83. McAllister, C.S., Taghavi, N., and Samuel, C.E. (2012). Protein kinase PKR amplification of interferon beta induction occurs through initiation factor eIF-2alpha-mediated translational control. J Biol Chem 287, 36384–36392. 10.1074/jbc.M112.390039.

84. Pierre, P., and Gatti, E. (2013). Loss of translation: a stealth weapon against pathogens? Nat Immunol 14, 1203–1205. 10.1038/ni.2759.

85. Dalet, A., Gatti, E., and Pierre, P. (2015). Integration of PKR-dependent translation inhibition with innate immunity is required for a coordinated anti-viral response. FEBS Lett 589, 1539–1545. 10.1016/j.febslet.2015.05.006.

86. Liang, Q., Deng, H., Sun, C.W., Townes, T.M., and Zhu, F. (2011). Negative regulation of IRF7 activation by activating transcription factor 4 suggests a cross-regulation between the IFN responses and the cellular integrated stress responses. J Immunol 186, 1001–1010. 10.4049/jimmunol.1002240.

87. Baruch, M., Hertzog, B.B., Ravins, M., Anand, A., Cheng, C.Y., Biswas, D., Tirosh, B., and Hanski, E. (2014). Induction of endoplasmic reticulum stress and unfolded protein response constitutes a pathogenic strategy of group A streptococcus. Front Cell Infect Microbiol 4, 105. 10.3389/fcimb.2014.00105.

88. Andreatta, M., and Carmona, S.J. (2021). UCell: Robust and scalable single-cell gene signature scoring. Comput Struct Biotechnol J 19, 3796–3798. 10.1016/j.csbj.2021.06.043.

89. Bolte, S., and Cordelieres, F.P. (2006). A guided tour into subcellular colocalization analysis in light microscopy. J Microsc 224, 213–232. 10.1111/j.1365-2818.2006.01706.x.

